# Type 2 diabetes Reprograms Bone Marrow Hematopoiesis and Dysregulates Immune Signaling in Response to Stroke

**DOI:** 10.64898/2025.12.29.696958

**Authors:** Hongxia Zhang, Wanjun Gu, Kailin Yu, Chia-Ling Tu, Wenhan Chang, Jialing Liu

**Affiliations:** Department of Neurological Surgery, University of California San Francisco, San Francisco, CA 94158; San Francisco VA Health Care System, San Francisco, CA 94158; Department of Neurology, University of California San Francisco, San Francisco, CA 94158; Department of Medicine, University of California San Francisco, San Francisco, CA 94158; University of California, Berkeley, CA

**Keywords:** diabetes, bone marrow, inflammation, stroke, hematopoiesis, scRNA-seq, immune dysregulation

## Abstract

Type 2 diabetes mellitus (T2DM) worsens stroke outcomes, but the underlying mechanisms linking T2DM to systemic immune dysfunction remain unclear. We investigated whether T2DM alters bone marrow (BM) hematopoiesis and dysregulate immune signaling following ischemic stroke in mice. Single-cell RNA sequencing, GeoMx digital spatial profiling (DSP), nCounter, and flow cytometry were used to analyze BM cells from control (db/+, Ctrl) and diabetic (db/db, T2DM) mice underwent experimental stroke or sham surgery. Diabetes caused marked structural remodeling of BM, with reduced cellularity and imbalance of hematopoietic lineages. Pseudotime trajectory analysis revealed impaired differentiation and maturation signatures of hematopoietic progenitor cells (HPC1) and granulocytes, and overactivation toward monocytes in diabetes after stroke. CellChat analysis demonstrated reorganization of intercellular communication, with hematopoietic progenitor cells (HPC1) and monocytes emerging as dominant signaling hubs through upregulated MIF, SIRP, and THBS pathways. AUCell enrichment indicated increased glycolysis and oxidative phosphorylation but reduced interferon-γ (IFN-γ) signaling, reflecting metabolic activation coupled with immune dysregulation. DSP and nCounter further confirmed upregulation of genes in the MIF, SIRP and THBS pathways in CD115⁺ monocytes and Ly6G⁺ neutrophils, indicating proinflammatory and migratory activation in diabetic bone marrow. Our data suggest that T2DM reprograms hematopoiesis and signaling networks, leading to maladaptive myeloid responses and impaired immune regulation after stroke. This maladaptive BM environment amplifies inflammation and limits repair, linking diabetic metabolic stress to worsened ischemic outcomes. Targeting bone marrow immune dysfunction may offer a therapeutic strategy to improve stroke recovery in diabetic patients.

## INTRODUCTION

Type 2 diabetes mellitus (T2DM) is a chronic metabolic disease associated with persistent inflammation, immune dysregulation, and increased risk of both cardiovascular and cerebrovascular complications^1^. Among these, ischemic stroke is particularly devastating. People with diabetes are not only more likely to suffer a stroke but also tend to experience greater injury and poorer functional recovery than those without diabetes^2,3^. While hyperglycemia, vascular dysfunction, and systemic inflammation clearly worsen cerebral ischemic damage, the upstream mechanisms linking diabetes to post-stroke immune response remain poorly defined.

As the primary site of hematopoiesis and major reservoir for various precursor cells, Bone marrow (BM) housed multipotent stem cell populations towards specific blood cell lineages^4^, including the hematopoietic stem cells (HSCs)^4,5^ that generate erythroid, myeloid, and lymphoid cells through intermediate progenitors^4,8,9^ to meet the varying and increasing demand of homeostatic and stress conditions^4,10,11^. In the steady-state, HSPCs also supply myeloid cells to tissues for immune surveillance and maintain reserves in the bone marrow that can be dispatched quickly when a threat is detected^4,6,7^. Under acute injury, the marrow activates emergency myelopoiesis^6,12^, rapidly expanding progenitors and producing neutrophils and monocytes required for debris clearance^13,14^, vascular remodeling^15–17^, and resolution of inflammation^13,18^. Diabetes disrupts this process by remodeling the hematopoietic niche^19,20^, reducing marrow cellularity^21^, biasing lineage output^6^, and sustaining inflammatory signaling^22^. Chronic metabolic stress alters stem and progenitor function^23^, suppresses lymphoid differentiation^24–26^, and skews myelopoiesis toward inflammatory states^23,25^. Whether and how these changes impair adaptive bone marrow responses to ischemic stroke remains unclear.

Like infection, ischemic stroke triggers a rapid, systemic immune response shaped by coordinated mobilization of bone marrow-derived cells^27^. In diabetic conditions, this response becomes exaggerated and poorly resolved^28,29^, contributing to blood-brain barrier (BBB) disruption, sustained secondary injury, and delayed recovery ^27,30^. It remains unclear whether this dysregulation originates from intrinsic reprogramming within the diabetic bone marrow or from downstream peripheral dysfunction cues. Because bone marrow–derived monocytes, neutrophils, and progenitors play key roles in post-stroke inflammation and repair, even subtle defects in their function can have profound effects on recovery. Diabetes, by chronically altering hematopoietic balance and immune tone, may therefore prime the bone marrow toward an exaggerated inflammatory response when ischemic injury occurs.

To address this, we integrated single-cell RNA sequencing (scRNA-seq) and spatial transcriptomics to define how type 2 diabetes alters immune cell populations and signaling in the bone marrow and validated the findings using immunohistology, flow cytometry and multiplex gene expression analyses. This integrated approach allowed direct assessment of hematopoietic development, metabolic pathway activity, and intercellular signaling networks at single-cell resolution. We tested the hypothesis that chronic metabolic stress in T2DM reprograms bone marrow hematopoiesis, disrupts emergency myelopoiesis following stroke, and drives maladaptive myeloid activation that worsens ischemic outcomes.

## MATERIALS AND METHODS

### Animals

C57BLKS/J mice carrying the leptin receptor mutation (B6.BKS(D)-*Lepr^db/db^*/J) were used as a model of obesity-induced type 2 diabetes mellitus (T2DM)^31^. Heterozygous littermates (db/+, B6.BKS(D)-*Lepr^db/+^*/J) served as normoglycemic controls. Genotyping was confirmed *via* toe clipping at postnatal day 7. Both male and female mice aged 6–10 months were included in all experiments. Animals were housed in groups of five under a 12-hour light/dark cycle with ad libitum access to food and water. All procedures were approved by the Institutional Animal Care and Use Committee (IACUC) at the San Francisco Veterans Affairs Medical Center and were conducted in accordance with NIH guidelines for the Care and Use of Laboratory Animals.

### Ischemic Stroke Model

The distal middle cerebral artery occlusion (dMCAO) model was performed based on previously established protocols^32,33^. Mice were anesthetized with 3% isoflurane for induction and maintained on 2% isoflurane in a 30% O₂/70% N₂O gas mixture via facemask. Core body temperature was maintained at 37□±□0.5□°C using a servo-controlled heating blanket and rectal probe. In the supine position, a midline neck incision was made, and the left common carotid artery (CCA) was exposed and ligated proximally to its bifurcation using 7-0 nylon suture. The mouse was then placed in the right lateral position. A 1-cm incision was made between the left eye and the ear (tragus). The temporal muscle was dissected to expose the skull, and the zygomatic arch was retracted to visualize the temporal bone. A 1-mm burr hole was drilled 2-mm rostral and superior to the foramen ovale to expose the proximal segment of the left MCA, which was then cauterized using bipolar forceps. After 60 minutes, the CCA ligature was removed. Sham-operated mice underwent the same procedure without occlusion of the MCA or CCA. Mice were sacrificed 3 days after stroke for further analysis.

### Bone Marrow Dissociation

Mice were anesthetized with isoflurane, and femurs and tibias were carefully dissected. Surrounding muscle tissue and periosteum were removed. Bones were cut at the joints, and bone marrow was flushed using a 23-gauge needle with 2 ml phosphate-buffered saline (PBS). The suspension was passed through a 70-μm cell strainer. Cells were collected by centrifugation, and red blood cells were lysed using 1× RBC lysis buffer (BioLegend, Cat# 420301) for 5 minutes at room temperature. All procedures were performed on ice unless otherwise specified.

### Single-Cell RNA Sequencing and Preprocessing

scRNA-seq was performed to characterize transcriptomic changes in bone marrow–derived cells under different experimental conditions, as previously described^34^. Bone marrow cells were isolated from mouse femurs and tibias, and single-cell suspensions were prepared as described above. Single cells were encapsulated into droplets using the 10x Genomics Chromium platform with GemCode technology, following the manufacturer’s instructions. Each cell and transcript were labeled with a unique cell barcode and molecular identifier (UMI), enabling precise identification of single-cell transcriptomes. cDNA libraries were generated using the 10x Genomics Single Cell 3′ Reagent Kit v2 and sequenced on an Illumina platform. Sequencing data were processed using the Cell Ranger Single Cell Software Suite (v1.3). Base call (BCL) files were converted to FASTQ format, and sequencing reads were aligned to the Mus musculus reference genome (mm10) using the STAR aligner. Gene count matrices were generated with the Cell Ranger count pipeline, and mean cluster transcript expression was analyzed with the Cell Ranger R kit. UMI counts were normalized by dividing each cell’s UMI count by its total UMI count and multiplying by the median of total UMI counts across all cells to ensure comparability between samples.

### Data Processing and Annotation

The gene count matrix was imported into Seurat (v5) for downstream analysis. Low-quality cells were filtered out based on the following criteria: total UMIs < 800, detected genes < 500, or mitochondrial read percentage > 10%. A total of 89,938 cells were retained in the final dataset. Batch effects were corrected using the FindIntegrationAnchors function. Expression data were log-normalized with a scaling factor of 10,000, the top 2,000 variable features were identified, the data were scaled, and principal component analysis (PCA) was performed. Differentially expressed genes (DEGs) for each cluster were identified using the FindAllMarkers function in Seurat (v5) with the Wilcoxon rank-sum test. Cell type annotation was performed using unsupervised clustering followed by reference-based cell type prediction using SingleR in combination with the Tabula Muris dataset (tissue: bone marrow). Annotated cell clusters were visualized using uniform manifold approximation and projection (UMAP). To visualize marker gene expression, the top 5-10 DEGs for each cluster were used to generate heatmaps, highlighting transcriptional differences among bone marrow cell populations. Dot plots were also generated for 2–5 representative markers from the top 10 DEGs per cluster. Feature plots display expression distribution in UMAP of representative gene marker of each cell cluster.

### Trajectory Analysis

Pseudotime trajectory analysis was performed using the Monocle 3 package to infer lineage relationships and assess developmental trajectories among bone marrow cell populations. Normalized gene expression data were extracted from the Seurat object and converted into Monocle format. Highly variable genes were selected to construct the trajectory graph and order cells along pseudotime, representing inferred developmental status. Dimensionality reduction was performed using uniform manifold approximation and projection (UMAP) for low-dimensional visualization of the developmental landscape. Pseudotime distributions were examined across experimental groups and within each annotated cell type to identify shifts in cellular states and lineage progression associated with diabetic and post-stroke conditions. This analysis provided a high-resolution framework to evaluate alterations in hematopoietic dynamics and differentiation pathways under metabolic and ischemic stress.

### Differentiation and Maturation Signature Analysis Using AUCell

AUCell quantifies gene set enrichment by calculating the Area Under the Curve (AUC) score for each gene set across the ranked expression values of all genes within a cell. This ranking prioritizes highly expressed genes, enabling cell-specific evaluation of gene set activity.

The defined gene sets in terms of differentiation and maturation in myeloid, monocytes and granulocytes were scored in single-cell RNA-seq data, with computing in downsized whole bone marrow (10,000 each group), 15,000 hematopoietic precursor cells (HPC1), 17,609 monocytes/promonocytes, and 13,121 granulocytes/granulocytopoietic cells across all four experimental groups. Statistical comparisons between groups were performed using Wilcox test.

### Cell–Cell Communication Analysis Using CellChat

Cell–cell communication analysis was performed using CellChat with default parameters^35^. Ligand–receptor interactions were inferred using curated databases and assessed for statistical significance using a permutation test with 100 permutations. Interactions with *p* < 0.05 were considered significant. Only cell populations containing at least 10 cells were included, and ligands and receptors were required to be expressed in at least 10% of cells in the corresponding populations.

Cell–cell communication networks were inferred using the CellChat R package (v1.6.1), which systematically analyzes ligand–receptor interactions based on single-cell transcriptomic data. The normalized expression matrix and cell type annotations from the Seurat object were used as inputs. A CellChat object was created using the createCellChat function, with the database set to CellChatDB.mouse for murine ligand–receptor interactions.

Lowly expressed genes were filtered using the subsetData function to reduce background noise. Interaction probabilities between cell populations were computed using computeCommunProb, aggregated with computeCommunProbPathway to identify enriched signaling pathways. The overall communication network was summarized using aggregateNet. Significant ligand–receptor pairs were identified using subsetCommunication to detect key signaling relationships between cell groups.

Network information from each group was merged to generate bubble plots illustrating ligand–receptor interactions for each cell cluster. Network centrality scores were calculated with netAnalysis_computeCentrality and visualized using scatter plots (netAnalysis_signalingRole_scatter) and heatmaps (netAnalysis_signalingRole_heatmap). Global communication patterns were visualized with chord diagrams using netVisual_aggregate and signaling gene expression distributions were displayed with violin plots generated by plotGeneExpression.

### Metabolism and Inflammation Signature Analysis Using AUCell

Pathway activity was analyzed using AUCell^36,37^ with the mouse Hallmark gene set (https://www.gsea-msigdb.org/gsea/msigdb/mouse/collections.jsp) to identify cells with active gene regulatory programs in single-cell RNA-seq data.

Hallmark signature scores were computed using AUCell for 15,000 hematopoietic progenitor cells (HPC1), 17,609 monocytes/promonocytes, and 13,121 granulocytes/granulocytopoietic cells across all four experimental groups. Analyses included metabolic pathways (glycolysis, adipogenesis, oxidative phosphorylation) and interferon pathway. Statistical comparisons between groups were performed using Wilcox test.

### GeoMx Digital Spatial Profiling (DSP) and Data Analysis

Formalin-fixed, paraffin-embedded (FFPE) femur bone sections (5 µm thick) were prepared from experimental mice for spatial transcriptomic analysis using the GeoMx® Digital Spatial Profiler (DSP, NanoString Technologies) according to the manufacturer’s protocol^38^. Sections were deparaffinized, rehydrated, and subjected to antigen retrieval in Tris-EDTA buffer (pH 9.0, 100°C for 15 minutes), followed by enzymatic digestion with proteinase K (0.1 µg/ml, 10 minutes at 37°C). Tissue sections were post-fixed with 10% neutral-buffered formalin (NBF) for 5 minutes, washed twice with NBF stop buffer (10 minutes each), then washed once with 1× PBS for 5 minutes. Sections were hybridized overnight with the NanoString mouse Whole Transcriptome Atlas (WTA) probe mix, followed by stringent washes with 50% formamide in 1× saline sodium citrate (SSC) buffer.

Bone sections were blocked with Buffer W for 1 hour at room temperature (RT) and incubated overnight at 4°C with primary antibodies against the monocyte marker CD115 (BioLegend, rabbit, 1:100) and the neutrophil marker Ly6G conjugated with Fluo-647 (BioLegend, rat, 1:100). After washing with 1× PBS, sections were incubated for 1 hour at RT with a secondary antibody (Fluo488-conjugated anti-rabbit, 1:400) and the nuclear stain SYTO83. Fluorescent images were acquired using the GeoMx DSP system to visualize tissue morphology and guide region of interest (ROI) selection. We defined candidate ROIs using CD115⁺ and Ly6G⁺ signals, then randomly sampled ROI from each signal set within each bone marrow section (Fig. 9A). Indexed oligonucleotide tags from each ROI were photocleaved using UV light and collected by microcapillary aspiration into a 96-well plate. The oligonucleotides were amplified by PCR, purified with AMPure XP beads, and quantified using Qubit. Amplified libraries were sequenced on an Illumina NovaSeq 6000 platform.

GeoMx RNA expression data were analyzed in R using the GeomxTools package for quality control, filtering, normalization, and dimensionality reduction with UMAP. The GeoMx object was then converted into a Seurat object for differential gene expression analysis, Gene Ontology (GO) enrichment, and gene expression visualization in CD115⁺ and Ly6G⁺ compartments separately.

### Gene Expression Analysis Using NanoString nCounter

Bone marrow cell suspensions were prepared as described above, and monocytes were isolated using the EasySep™ Mouse Monocyte Isolation Kit (STEMCELL Technologies, Cat# 19861) according to the manufacturer’s protocol. Cell suspensions were incubated with the EasySep™ monocyte enrichment cocktail and magnetic nanoparticles. Unwanted cells were magnetically labeled and removed using the EasySep™ magnet, yielding an untouched population of highly enriched monocytes.

Isolated monocytes were lysed in TRIzol reagent (Life Technologies, Cat# 15596018), and RNA was extracted using the phenol-chloroform method followed by cleanup with the PureLink™ RNA Mini Kit (Thermo Fisher Scientific, Cat# 12183018A). RNA quality and concentration were assessed using the Agilent RNA 6000 Pico Kit (Agilent, Cat# 5067-1513) on a Bioanalyzer 2100 system (Agilent).

Gene expression profiling was performed using the NanoString nCounter® Myeloid Innate Immunity Panel v2. GeoMx DSP counts were normalized using the geometric mean of housekeeping genes to account for technical variability across regions of interest and groups. Data normalization and pathway analysis were conducted using nSolver 4.0 software (NanoString Technologies, Seattle, WA, USA) according to the manufacturer’s recommendations. Gene expression boxplots were generated in R.

### Fluorescence-Activated Cell Sorting (FACS)

To quantify lineage-specific cell populations in bone marrow, single-cell suspensions were prepared after red blood cell lysis as described above. Cells were washed and resuspended in staining buffer (0.5% BSA, 2 mM EDTA in 1× PBS) and incubated with Fc Block (BioLegend) for 20 minutes on ice to minimize nonspecific binding. Cells were then stained on ice for 30 minutes using three antibody panels:

a) **Precursor cells:** Biotin anti-mouse Lineage Panel (anti-CD3ε [145-2C11], anti-Ly6G/Ly6C [RB6-8C5], anti-CD11b [M1/70], anti-CD45R/B220 [RA3-6B2], anti-TER-119 [TER-119]; BioLegend, 133307), biotin anti-CD4 (GK1.5, 100403, BioLegend), biotin anti-CD8a (53-6.7, 100703, BioLegend), biotin anti-CD11c (N418, 117303, BioLegend), biotin anti-NK1.1 (PK136, 108703, BioLegend), c-Kit BV605 (2B8, 105847, BioLegend), Sca-1 APC-Cy7 (D7, 108126, BioLegend), CD150 PerCP-Cy5.5 (TC15-12F12.2, 108126, BioLegend), CD48 Pacific Blue (HM48-1, 103418, BioLegend), CD34 AF700 (RAM34, 56-0341-82, eBioscience™), CD16/32 PE-Cy5 (S17011E, 156618, BioLegend), and CD115 APC (AFS98, 135510, BioLegend).
b) **Myeloid cells:** CD3 FITC (17A2, 11-0032-82, eBioscience™), B220 eFluor450 (RA3-6B2, 48-0452-82, eBioscience™), CD11b PerCP-Cy5.5 (M1/70, 45-0112-82, eBioscience™), Ly6G APC-eFluor780 (1A8, 127624, BioLegend), CD115 APC (AFS98, 135510, BioLegend), Ly6C BV605 (HK1.4, 128036, BioLegend), and CD11c PE-Cy7 (N418, 117318, BioLegend).
c) **Lymphocytes:** Biotin anti-mouse Lineage Panel, c-Kit BV605 (2B8, 105847, BioLegend), Sca-1 APC-Cy7 (D7, 108126, BioLegend), IL7Rα PerCP-Cy5.5 (SB/199, 121114, BioLegend), B220 eFluor450 (RA3-6B2, 48-0452-82, eBioscience™), and CD93 APC (AA4.1, 136510, BioLegend).

Biotin-conjugated antibodies were detected using FITC Streptavidin (405202, BioLegend) incubated for 30 minutes on ice. Dead cells were excluded using Fixable Viability Dye eFluor506 (65-0866-14, eBioscience™). Stained cells were analyzed on a BD FACS Fusion 3 flow cytometer, and data were processed using FlowJo v10.8.

### Immunofluorescence staining (IF)

Immunofluorescence staining was performed on bone marrow sections as previously described^39^. The primary antibodies used were as follows: CD74 (1:200, rat, BioLegend) and CD115 (1:100, rabbit, BioLegend). The secondary antibodies used were as follows: Alexa Fluor 488-, 594-conjugated donkey anti-rat, anti-rabbit IgG (1:400: Jackson Laboratory). Fluorescence signals were detected with confocal scanning microscope.

### Hematoxylin and eosin (H&E) staining

Formalin-fixed, paraffin-embedded (FFPE) femur bone sections from all four experimental groups were stained with hematoxylin and eosin stain (H&E), as preciously described^40^. In brief, the sections were deparaffinized and rehydrated, stained in hematoxylin for 10 minutes, rinsed in tap water, differentiated in 1% acid alcohol for a few seconds, rinsed in tap water, stained Eosin Y for 2 minutes, and then dehydrated and cleared and mounted with permanent mounting medium.

### Statistical analysis

All Flow cytometry data were analyzed using GraphPad Prism (v8.0). Data is presented as mean ± standard error of the mean (SEM) unless otherwise stated. Normality was assessed using one of these tests: Anderson-Darling, D’Agostino-Pearson omnibus, Shapiro–Wilk and Kolmogorov-Smirnov test. For comparisons between two groups, two-sided t-test was applied. For multiple group comparisons, two-way ANOVA followed by Tukey’s post hoc test was used as appropriate.

Transcriptomic data were analyzed using R (v4.3.0). Differential gene expression analyses in single-cell RNA sequencing were performed using the FindAllMarkers function in Seurat (v5) with the Wilcoxon rank-sum test. P values were adjusted for multiple comparisons using the Bonferroni or Benjamini–Hochberg correction as indicated. Pseudotime, cell–cell communication, and pathway enrichment analyses were conducted using Monocle 3, CellChat (v1.6.1), and AUCell, respectively, following each package’s standard workflow.

NanoString nCounter gene expression data were normalized using nSolver 4.0 software according to the manufacturer’s protocol. Spatial transcriptomic data from the GeoMx DSP platform were processed and normalized using GeomxTools in R, and differential gene expression and Gene Ontology (GO) enrichment were analyzed after conversion to Seurat objects. For all analyses, an adjusted p value less than 0.05 was considered statistically significant.

## RESULTS

### Single-Cell Transcriptomic Map of Bone Marrow Across Diabetic and Stroke Conditions

To generate a reference map of hematopoietic populations in the BM, BM cells were collected and pooled from both femurs and tibias of three mice per experimental group. Following RBC lysis, single-cell RNA sequencing (scRNA-seq) was performed on BM single cells from four groups: control (Ctrl; db/+), type 2 diabetes mellitus (T2DM; db/db), stroke (Stroke; db/+ Stroke), and T2DM with stroke (T2DM.Stroke; db/db Stroke). Following quality control and filtering, a total of 89,938 high-quality cells were retained for downstream analysis. Quality metrics confirm consistent RNA integrity and sequencing depth across all samples, as reflected by comparable numbers of detected genes (nFeature_RNA), total unique molecular identifiers (nCount_RNA), and low mitochondrial transcript proportions (percent.mt). Median per-cell metrics included 824 unique genes and 2,063 unique molecular identifiers (UMIs), with minimal mitochondrial read content, confirming data quality and cross-sample comparability (data not shown). Dimensionality reduction and visualization were performed using Uniform Manifold Approximation and Projection (UMAP). Nine distinct clusters were resolved through an integrated framework combining reference-based annotation, transcriptional similarity, and inferred developmental positioning (Figure 1A). The spatial distribution of these clusters reflected established differentiation hierarchies within the BM microenvironment. Hematopoietic progenitor populations were separated into distinct clusters representing hematopoietic progenitor cells (HPC1), megakaryocyte/erythroid precursors (HPC2), and pro-pre-B cells (HPC3) based on progenitor features and lineage bias. Granulocytopoietic cells were distinguished from mature granulocytes based on transcriptional maturity and developmental state, allowing separation of immature and terminal granulocytic compartments. Promonocytes were resolved as an intermediate population between progenitors and monocytes, reflecting transitional monocytic differentiation states. Monocytes were distinguished from macrophage-like predictions based on bone marrow localization and lack of tissue-resident macrophage characteristics. Basophils and proerythroblasts formed discrete, well-resolved clusters with strong concordance between clustering and reference annotation. Ambiguous or overlapping reference assignments were resolved by prioritizing lineage consistency, developmental trajectory positioning, and population-level transcriptional coherence, ensuring that each cluster was assigned a single biologically coherent identity. Raw cell counts for each resolved cell type were extracted from the full dataset and normalized across experimental groups prior to proportional comparisons, enabling robust assessment of hematopoietic composition changes across conditions. The high concordance of annotation between Seurat clusters and reference displayed in supplementary Table 1. The canonical lineage markers identified for each populations: hematopoietic progenitor cells (HPC1; Elane, Ctsg, Prtn3), granulocytopoietic cells (Ltf, Lcn2, Camp), promonocytes (Ly6c2, Fn1, Lyz2, Crip1), monocytes (Fn1, Lyz2, Crip1), granulocytes (Ltf, Lcn2, Ngp, Camp, Ifitm6), megakaryocyte/erythroid precursors (HPC2; Car2, Car1, Prdx2, Hba-a1), proerythroblasts (Car2, Car1, Prdx2), basophils (Prss34, Mcpt8), and pro-pre-B cells (HPC3; Cd74, Cd79a) (Figure 1B). UMAP feature plots show the spatial distribution and relative expression of selected marker genes across all nine identified clusters (Supplementary Figure 1A). A heatmap of top up to 10 differentially expressed genes in each major population further confirmed distinct lineage-specific transcriptional programs across the identified clusters (Supplementary Figure 1B). Collectively, this high-resolution cellular transcriptomic dataset in the bone marrow establishes a foundational framework for subsequent comparative analyses of hematopoietic dynamics across experimental conditions.

**Figure 1.**
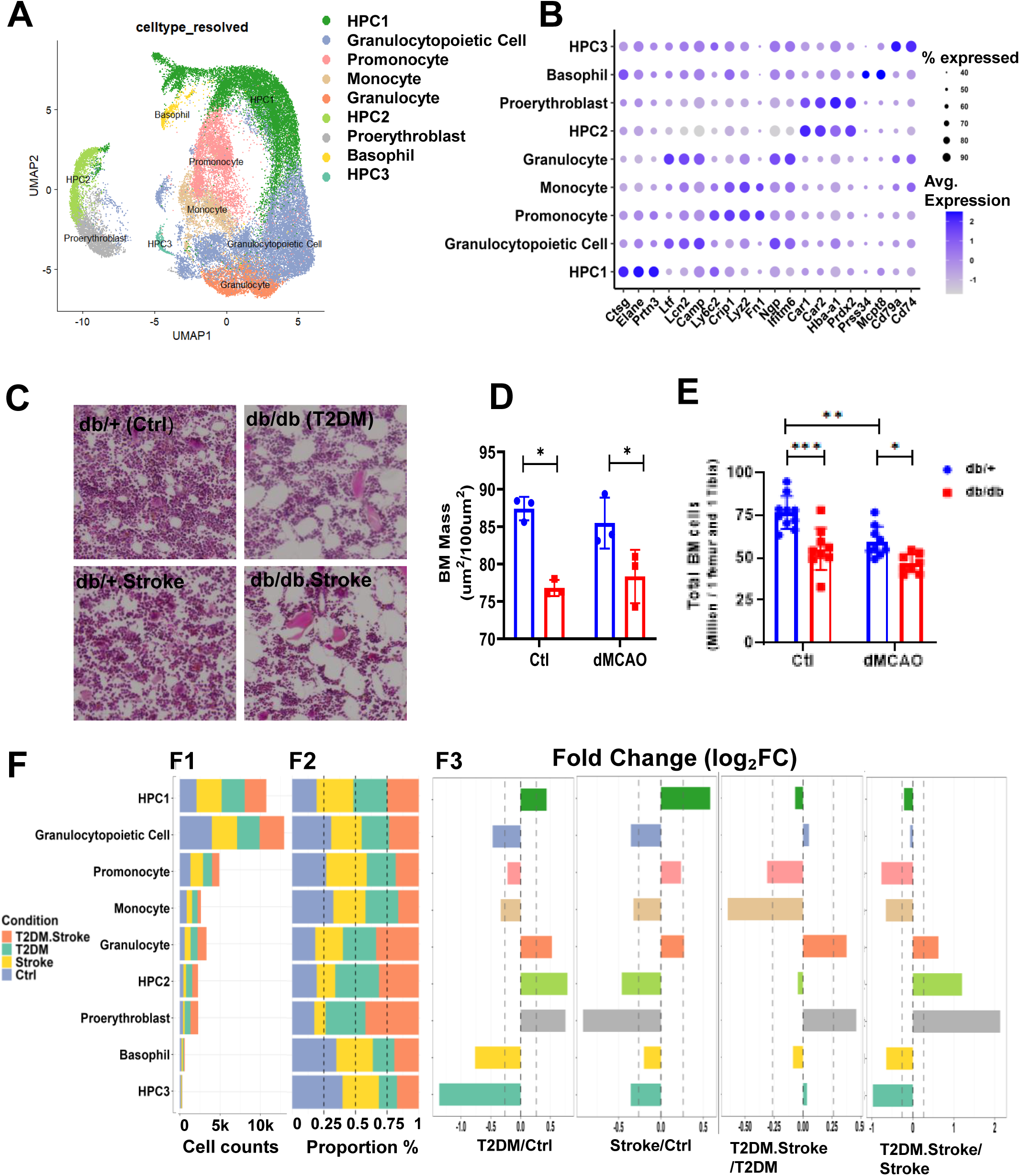
A comprehensive scRNA-seq map and diabetes- and stroke-associated alterations in bone marrow cellular and composition. **(A)** Uniform Manifold Approximation and Projection (UMAP) visualization of the integrated bone marrow single-cell RNA-sequencing dataset across all four experimental groups (Ctrl, T2DM, Stroke, and T2DM + Stroke) following batch correction. Cell clusters were identified by unsupervised Seurat clustering and annotated using SingleR in combination with the Tabula Muris bone marrow reference dataset. **(B)** Dot plot showing scaled average expression (color intensity) and percentage of expressing cells (dot size) for representative lineage markers used for cluster annotation. This panel illustrates transcriptional distinctions among hematopoietic progenitors, granulocytic, monocytic, erythroid, basophil, and lymphoid compartments (full marker sets shown in Supplementary Figure 1 and Table 1). **(C)** Representative hematoxylin and eosin (H&E)–stained femoral bone marrow sections from control and diabetic mice with or without ischemic stroke, illustrating changes in marrow cellularity and adiposity. Images were acquired from comparable femoral regions. **(D)** Quantification of bone marrow cellular mass expressed as marrow cellular area fraction (µm² per 100 µm² total marrow area). Diabetic mice exhibit reduced marrow cellular area compared with controls, independent of stroke. N=3. **(E)** Total nucleated bone marrow cell counts measured from one femur and one tibia per mouse after red blood cell lysis at the indicated post-stroke time point. Each dot represents an individual mouse; bars show mean ± SEM. N= 8-10. **(F)** Quantitative analysis of bone marrow cell-type composition derived from scRNA-seq data following random down-sampling to 10,000 cells per group. F1: Absolute cell numbers per annotated cluster by condition after down-sampling. F2: Within-group cell-type composition, shown as the fraction of each cluster relative to total cells in that experimental group. (e.g., number of HPC1 cells in control group / total cells in control group, grouped by all groups). Coded color represents each group: Ctrl, Stroke, T2DM and T2DM.Stroke. F3: Log₂ fold change (log₂FC) in average cell-type fraction for selected comparisons (T2DM vs Ctrl, Stroke vs Ctrl, T2DM + Stroke vs T2DM, and T2DM + Stroke vs Stroke). Dashed lines at ±1.2 log₂FC indicate visualization cutoffs for highlighting large compositional shifts. Coded color represents each cell cluster. For all panels, Ctrl = db/+ sham; T2DM = db/db sham; Stroke = db/+ dMCAO; T2DM + Stroke = db/db dMCAO. Statistical significance is indicated as follows: \**P* < 0.05; \*\**P* < 0.01; \*\*\**P* < 0.001.

### Diabetes and Stroke Impair Bone Marrow Cellularity and Reshape Hematopoietic Composition

To assess the impact of diabetes and stroke on bone marrow architecture and hematopoietic composition, we integrated histological staining, cell enumeration, single-cell transcriptomics, and flow cytometry. Hematoxylin and eosin (H&E) staining of femoral BM sections revealed pronounced structural alterations in db/db (T2DM) mice compared with db/+ (control) animals. Diabetic BM exhibited reduced nucleated cell density and increased adipocyte infiltration (Figure 1C and Figure 8A). Quantitative morphometric analysis confirmed a significant reduction in marrow mass areas in diabetic groups, regardless of stroke status, compared with non-diabetic groups(Figure 1D).

To confirm these histological findings, total BM cellularity was quantified after red blood cell (RBC) lysis using both hemocytometer counting and flow cytometry with counting beads. Diabetic mice (db/db) showed a marked reduction in total BM cells relative to non-diabetic controls (db/+) under both baseline and post-stroke conditions (Figure 1E). Stroke decreased BM cellularity in db/+ mice, while the decrease in db/db mice was not significant (db/+: 76.5 ± 4.38 million; db/db: 55.2 ± 4.52 million; db/+ stroke: 59.6 ± 4.63 million; db/db stroke: 46.7 ± 4.49 million).

Single-cell RNA sequencing further delineated diabetes- and stroke-associated alterations in hematopoietic composition. To control for unequal cell numbers across groups, we randomly down-sampled to 10,000 cells in each group, HPC1, monocytes (including clusters of promonocyte and monocyte) and granulocytes (including clusters of granulocytopoietic cell and granulocyte) comprise the predominant cell populations in the bone marrow, collectively accounting for 86.2% of total cells (Figure 1F, F1). Cluster-level analysis demonstrated shifts in relative proportions across major hematopoietic lineages (Figure 1F, F2 and F3). Diabetic non-stroke (T2DM) and control stroke (Stroke) groups exhibited increased frequencies of HPC1 and granulocytes, compared to control non-stroke (Ctrl) group. They also exhibited decreased frequency of granulocytopoietic cells, monocytes, basophils, and pro-pre-B cells (HPC3), but increased megakaryocyte/erythroid precursors (HPC2) and proerythroblasts compared to both groups of non-diabetic controls, indicating a lineage bias toward megakaryocyte/erythroid differentiation in diabetes.

Comparison of cluster proportions (Figure 1F, F3) revealed that both stroke and diabetes increased HPC1 representation relative to their respective control, suggesting enhanced progenitor expansion under disease conditions. Promonocyte proportions decreased in diabetic mice after stroke (T2DM.Stroke) compared to diabetic mice (T2DM) and non-diabetic mice after stroke (Stroke), with monocytes further reduced in across all comparisons, indicating diabetes-exacerbated impairment of monocyte replenishment after stroke. Both diabetes and stroke reduced the frequencies of basophil and pro-pre-B cell (HPC3), while opposing effects were found on megakaryocyte/erythroid precursors and proerythroblasts.

Flow cytometric validation corroborated these compositional shifts. The gating strategy for hematopoietic stem cells (HSCs), progenitors, lymphoid cells, and myeloid lineages is shown in Supplementary Figure 2A-C. Diabetic mice with stroke increased proportions of Lineage⁻Sca-1⁺c-Kit⁺(LSK) stem/progenitor cells, compared to non-diabetic stroke mice. Granulocyte-monocyte progenitors (GMPs) were elevated in diabetic mice compared to non-diabetic mice under baseline conditions (Figure 2A-B). HSCs and monocyte-dendritic progenitors (MDPs) did not differ significantly across groups (Supplementary Figure 2D), suggesting preserved early myeloid precursor production. However, common lymphoid progenitors (CLPs) were markedly reduced after stroke in both db/+ and db/db mice (Figure 2C), and diabetic mice exhibited lower baseline CLP counts compared with non-diabetic controls (Supplementary Figure 2E). These reductions were accompanied by a pronounced decrease in pre-pro-B cells, particularly in the diabetic stroke group (Figure 2C and Supplementary Figure 2E), indicating diabetes-induced suppression of early lymphoid differentiation.

**Figure 2.**
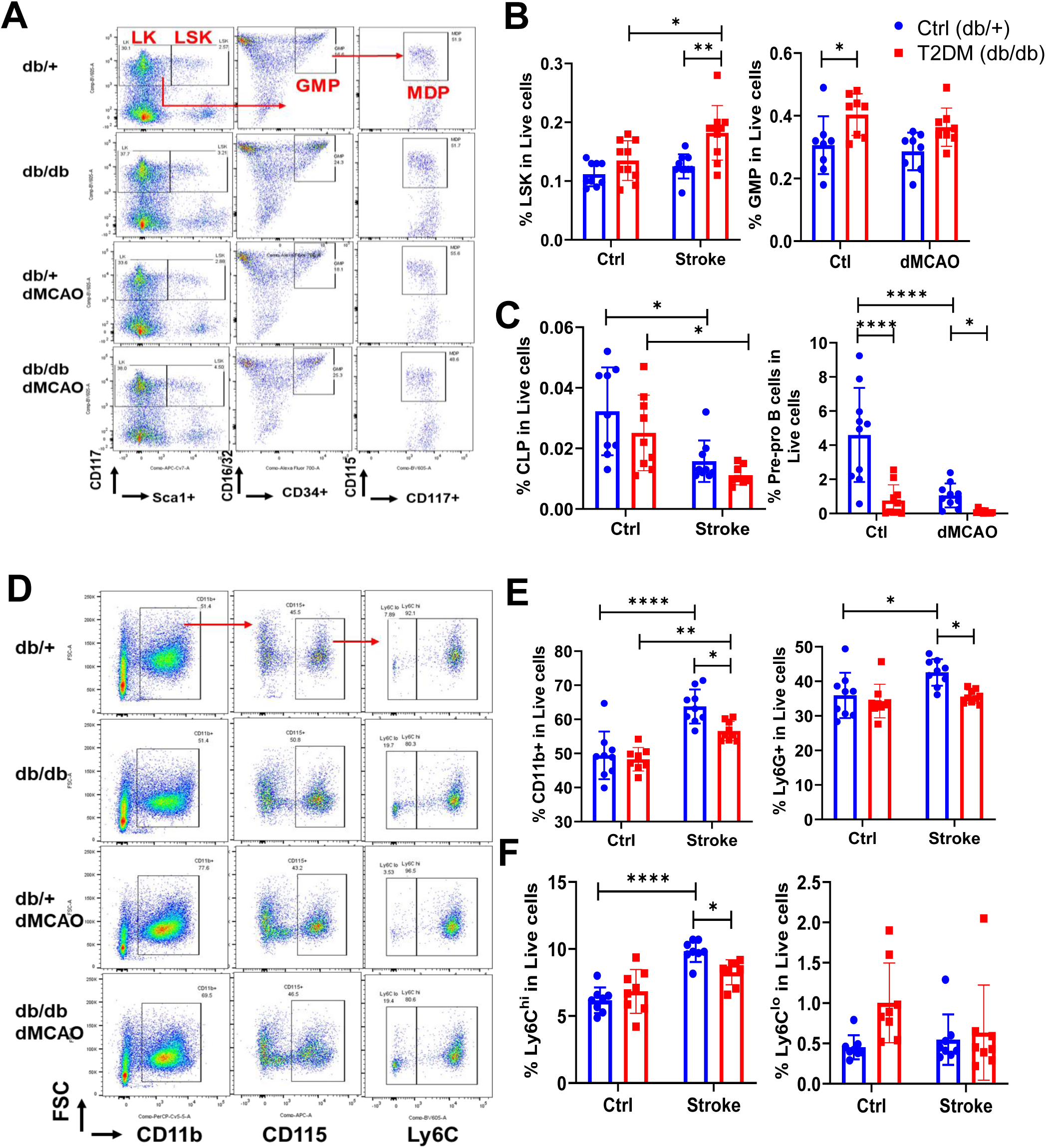
Flow cytometry (FACS) showing alterations in bone marrow hematopoietic lineages following stroke in control and diabetic mice. **(A)** Representative flow cytometry gating images of stem/precursors in the bone marrow across groups. **(B)** Flow cytometry (FACS) analysis showing altered proportions of hematopoietic stem and progenitor cells (LSK: Lineage⁻Sca-1⁺c-Kit⁺) and granulocyte–monocyte progenitors (GMPs) in live cells across the four groups. **(C)** FACS analysis of proportion changes of common lymphoid progenitors (CLPs) and pre-pro B cells in live cells across the four groups. **(D)** Representative flow cytometry gating images of myeloids in the bone marrow across groups. **(E)** FACS analysis of myeloid cells (CD45⁺CD11b⁺) and neutrophils (Ly6G⁺) across the four groups. **(F)** FACS analysis of monocytes, including Ly6C^high^ and Ly6C^low^ subsets, across the four experimental groups (Ctrl, Stroke, T2DM, and T2DM + Stroke). N= 8-10. Two-way ANOVA followed by Tukey’s post hoc test was used for analysis. Statistical significance is indicated as follows: \**P* < 0.05; \*\**P* < 0.01; \*\*\**P* < 0.001, \*\*\*\**P* < 0.0001.

Myeloid analysis revealed that total myeloid cells (CD11b⁺), neutrophils (Ly6G⁺), and Ly6C^hi^ monocytes were elevated in db/+ mice after stroke (Figure 2D-F). Diabetes increased total myeloid cells, but not neutrophils and Ly6C^hi^ monocytes after stroke in the bone marrow (Figure 2D-F). All myeloid subsets were significantly reduced in diabetic mice compared with non-diabetic mice after stroke (Figure 2D-F), while Ly6C^lo^ monocytes remained unchanged across groups (Figure 2F).

Collectively, these findings demonstrate that diabetes induces profound BM atrophy, suppresses lymphoid progenitor output, and skews hematopoietic differentiation toward the megakaryocyte/erythroid lineage at the expense of monocytes, leading to dysregulated myeloid dynamics within the diabetic BM niche.

### Diabetic Stroke Alters Bone Marrow Hematopoietic Development Trajectories

To investigate how stroke, type 2 diabetes mellitus (T2DM), and their comorbidity influence hematopoietic development, we performed pseudotime trajectory analysis of single-cell RNA sequencing data using Monocle 3. The UMAP visualization colored by pseudotime values revealed a continuous developmental trajectory, representing the inferred direction of progression from early progenitor populations toward late mature myeloid lineages (Figure 3A).

**Figure 3.**
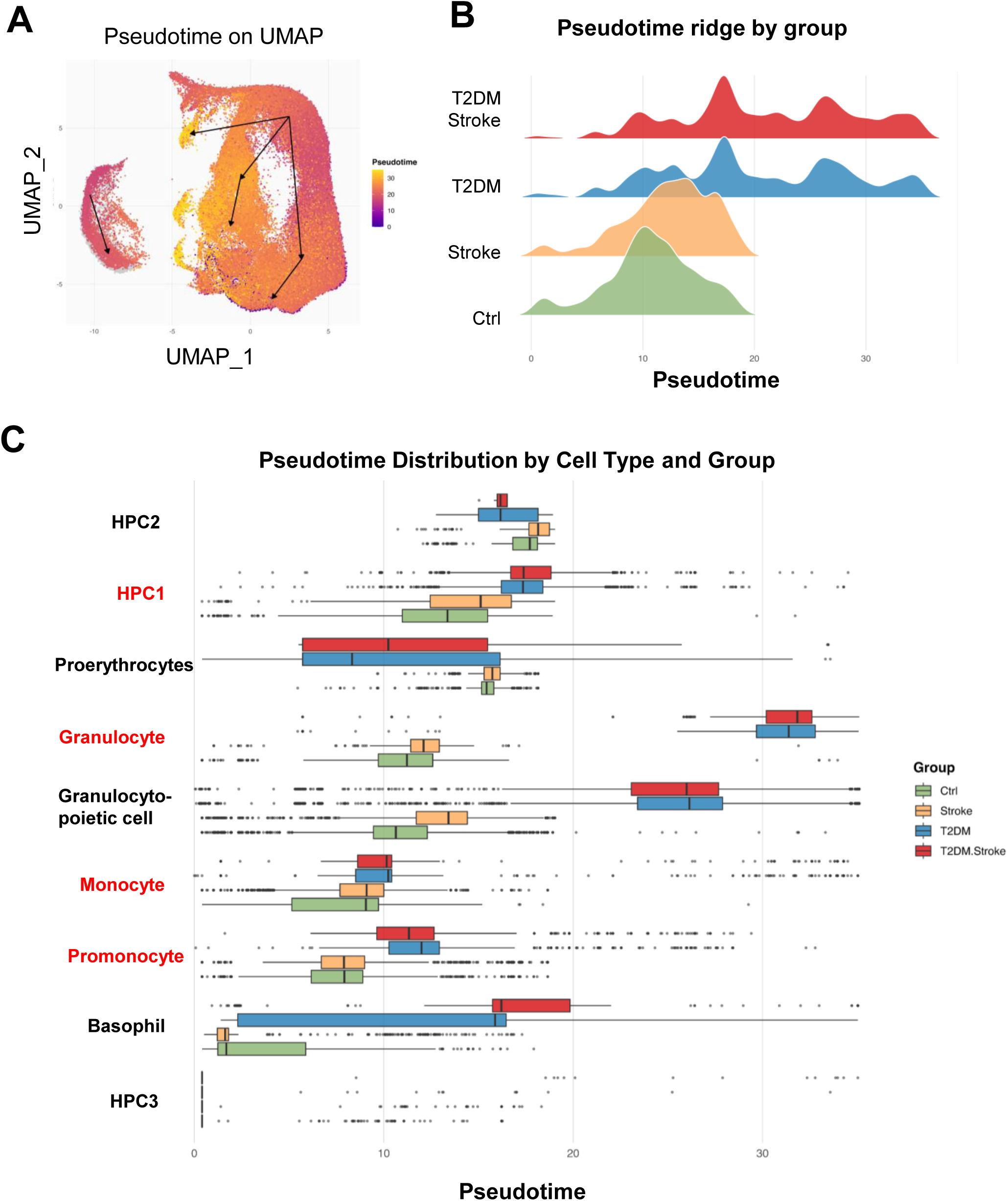
Trajectory analysis reveals developmental imbalance in diabetic stroke bone marrow. **(A)** UMAP visualization of bone marrow cell clusters colored by pseudotime values, illustrating directional cell state progression. Darker colors represent early status and lighter colors represent later status. Arrows and connecting lines indicate the inferred trajectory and direction of cellular progression across clusters in the bone marrow. **(B)** Ridge density plots show pseudotime distributions of cells across experimental groups (Ctrl, Stroke, T2DM, and T2DM + Stroke), revealing greater heterogeneity in the diabetic groups, with representation of both immature and mature populations. The x-axis represents pseudotime, reflecting progression along the transcriptional trajectory, while the y-axis indicates kernel density for each group. Peak height corresponds to cell frequency. **(C)** Boxplots showing pseudotime distributions for each cell cluster across conditions experimental groups (Ctrl, Stroke, T2DM, and T2DM + Stroke). Boxes indicate the interquartile range with median pseudotime values, and dots correspond to individual cells. This visualization enables comparison of trajectory positioning and heterogeneity across both cell types and groups. HPC1, granulocyte, granulocytopoietic cell, monocyte, promonocyte and basophil showed a shift toward later pseudotime values in T2DM and T2DM+Stroke conditions, indicating progression toward more developed or activated transcription cellular status.

Comparison of pseudotime distributions across the four experimental groups revealed distinct developmental dynamics (Figure 3B). Non-diabetic groups (Ctrl and Stroke) exhibited a prominent peak at early pseudotime values and simple model distribution, consistent with a more basal or less activated transcriptional state and predominance of cells at similar differentiation states, while the diabetic groups (T2DM and T2DM.Stroke) showed rightward shift toward later pseudotime and a broader distribution, suggesting a more developed or activated cellular status that reflects an increased cellular heterogeneity on the developmental scale.

Pseudotime distributions stratified by cell type and groups further delineated lineage-specific alterations (Figure 3C). Granulocytes, and granulocytopoietic cells in diabetic groups and basophils in diabetic stroke group showed a significant shift toward later pseudotime values in T2DM and T2DM+Stroke conditions, indicating progression toward more activated transcription cellular status. HPC1 and promonocytes in the diabetic groups have a slight increase in psuedotime values compared to the non-diabetic groups for the same tendency. These findings indicate that metabolic and ischemic stressors differentially remodel cellular trajectory dynamics, with the most pronounced effects observed in precursors, promonocytes and granulocytes lineages.

To further evaluate functional remodeling of bone marrow myeloid populations, we quantified predefined gene sets scores related to lineage differentiation that represents from stem/precursors to early lineage commitment^41,42^, and related to maturation that reflects transition from late stages to functionally readiness^5,43^. We calculated scores using AUCell function in R for myeloid differentiation and maturation in BM and HPC1, monocyte differentiation and maturation in monocyte and promonocyte, granulocyte differentiation and maturation in granulocyte and granulocytopoietic cell. Stroke induced a robust increase in differentiation signature across BM, HPC1 and granulocytes in non-diabetic mice. However, stroke decreased differentiation in all 4 cell types in diabetic ones (Figure 4A-B), suggesting a complex interplay between stroke and diabetes in the myeloid cell types. Diabetes also increased differentiation potential in monocytes in the absence of stroke.

**Figure 4.**
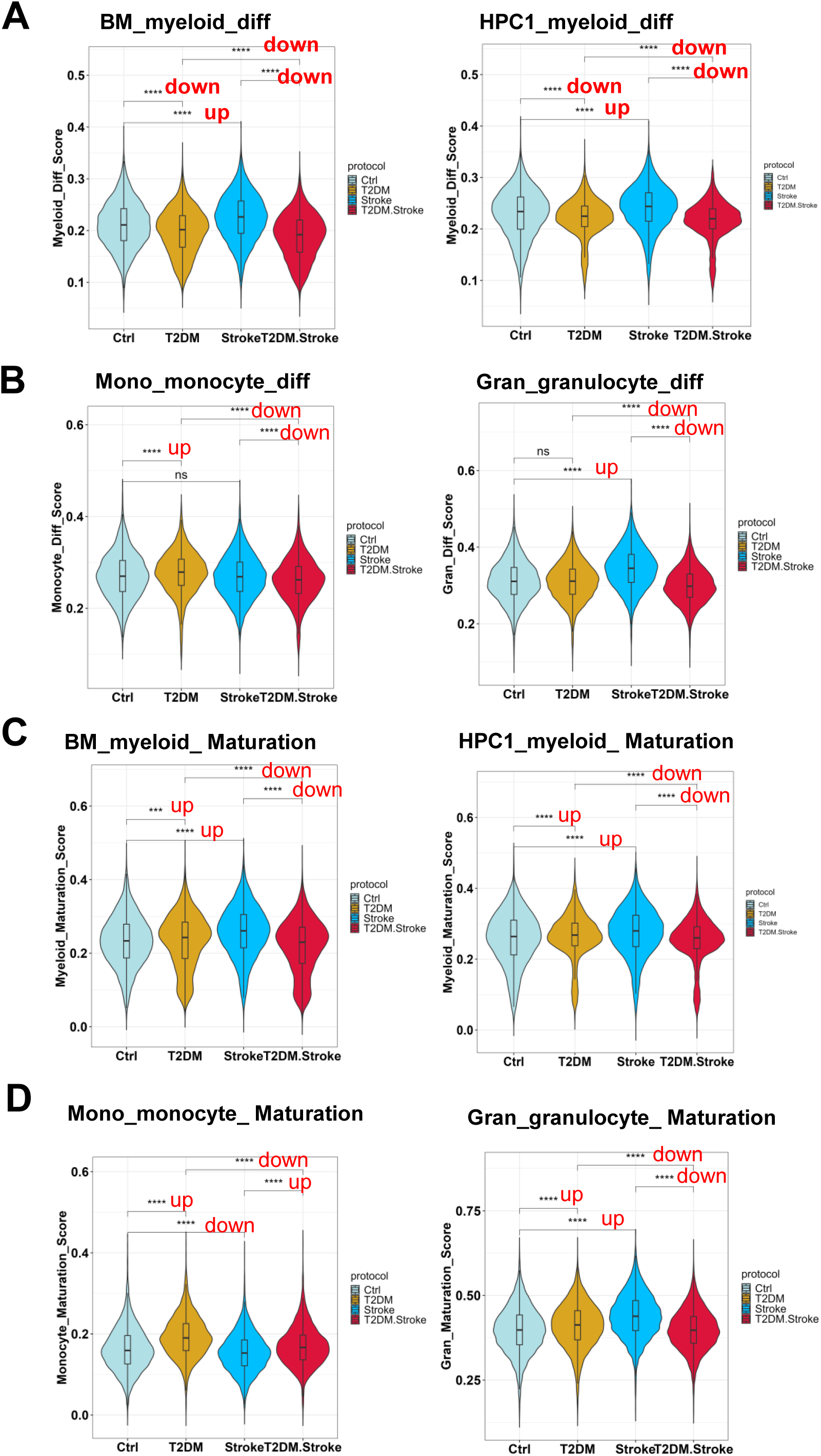
AUCell calculates differentiation and maturation signatures score. **(A)** Myeloid differentiation score of genes in whole bone marrow and HPC1. **(B)** Monocyte and granulocyte differentiation score of genes in monocytes and granulocytes. **(C)** Myeloid maturation score of genes in whole bone marrow and HPC1. **(D)** Monocyte and granulocyte maturation score of genes in monocytes and granulocytes. 10,000 down-sampled whole BM cells, 15,000 HPC1, 17,609 monocytes/promonocytes, and 13,121 granulocytes/granulocytopoietic cells across all four experimental groups were used for AUCell analysis. Statistical comparisons between groups were performed using Wilcox test, with asterisks indicating statistical significance.

As for maturation signature, the similar opposing effect of stroke in non-diabetic vs diabetic mice could be seen in BM, HPC1 and granulocytes (Figure 4C-D). Diabetes increased the maturation index in the monocytes of both non-stroke and stroke mice, consistent with a more developed or activated status. Together, these findings suggest that there appears to be a complex interplay between stroke and diabetes in activation and differentiation among the myeloid cell types in the marrow.

### Diabetic Stroke Alters Cell–Cell Communication Networks in Bone Marrow

To characterize changes in intercellular signaling within the bone marrow under pathological conditions, we performed CellChat network analysis to evaluate outgoing (signal-sending) and incoming (signal-receiving) interactions among hematopoietic cell populations. As shown in Figure 5A, both signaling output and input strengths (referring to total counts of statistically significant ligand–receptor pairs) varied across experimental groups. Besides HPC2, granulocytes maintained a relatively strong signal receiving cell type in the both of the non-diabetic groups, while monocytes and promonocytes emerging as the dominant signaling sources in the 3 pathological groups. These data suggest that the disease conditions reorganized intercellular communication hierarchies.

**Figure 5.**
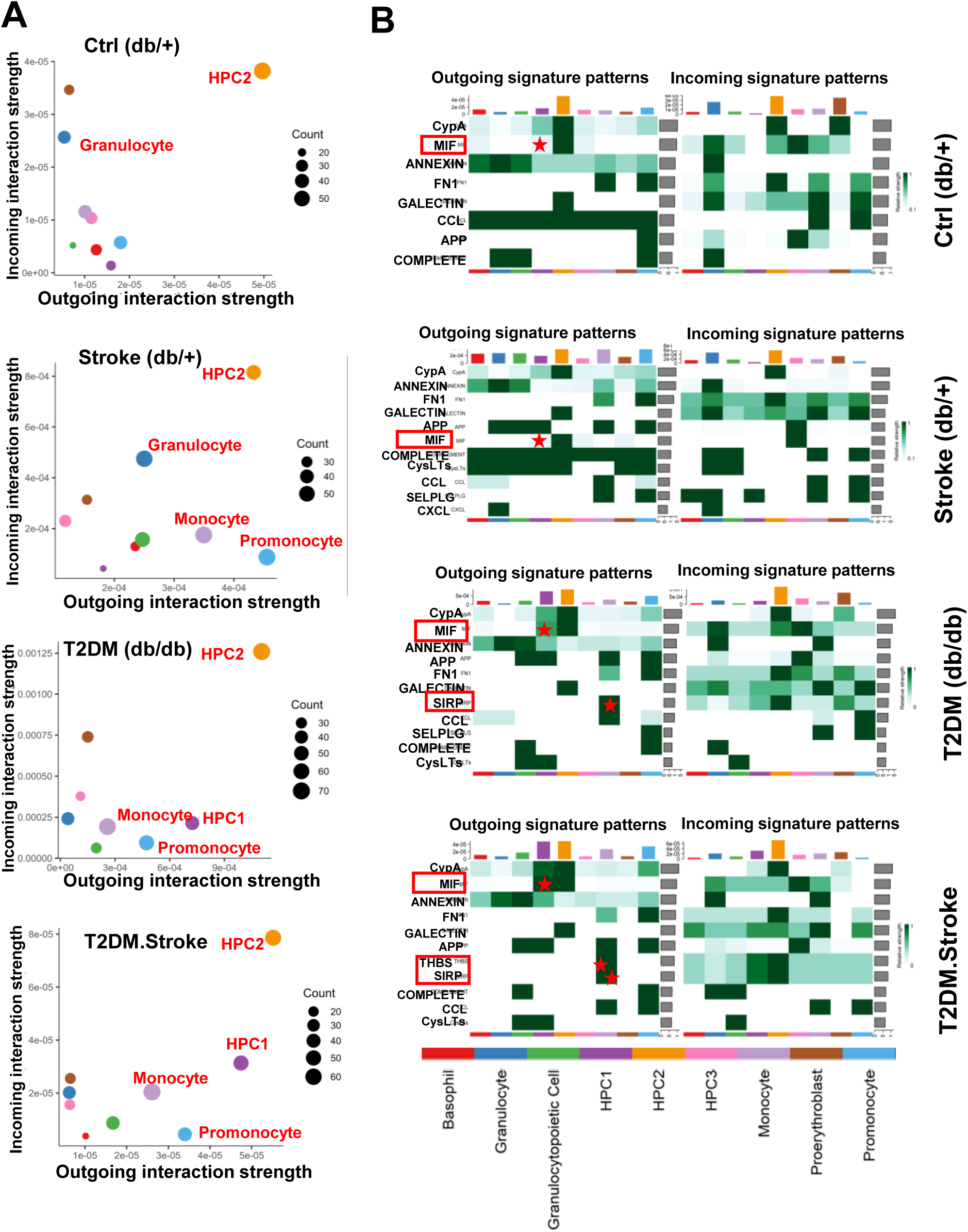
Cell–cell communication networks are altered in diabetic stroke bone marrow. **(A)** Scatter plots showing visualization of dominant signal-sending (outgoing interactions) and signal-receiving (incoming interactions) cell clusters based on CellChat network analysis. Each cluster is plotted according to its total outgoing and incoming signaling strength (defined by total counts of statistically significant ligand–receptor pairs), with point size representing cell numbers of statistically significant ligand-receptor pairs per group. HPC1, monocytes, and promonocytes emerge as the primary signal senders in the Stroke, T2DM, and T2DM + stroke groups. **(B)** Heatmaps showing predicted significant intercellular signaling between annotated bone marrow clusters across experimental conditions. For each group, the left panel displays outgoing (ligand-enriched) signaling, and the right panel shows incoming (receptor-enriched) signaling. The displayed signaling in each group is defined by the computed interaction probability of specific ligand-receptor pairs mediated between a sender and a receiver cell population. Signaling strength, based on CellChat interaction probability, is row-scaled by a specific signaling pathway and represented by a color gradient (color scale shown at right). Bar plots adjacent to each heatmap summarize the total signaling strength across all pathways. Red frames highlight increased outgoing signaling of MIF, SIRP, and THBS pathways in the T2DM and T2DM + stroke groups. CellChat analysis was performed with default parameters displayed in method. Only statistically significant interactions identified by CellChat are displayed.

The heatmaps in Figure 5B display the predicted intercellular signaling based on the interaction probability of ligand-receptors and relative strength between clusters. Diabetes significantly elevated outgoing signaling activity through the MIF, SIRP, and THBS pathways (Figure 5B, highlighted in red frames), which were active predominantly in signal-sending populations such as HPC1 and monocytes as highlighted in Figure 5B (red asterisks), implicating them in inflammatory and stress-related signaling cascades. Collectively, these results demonstrate that stroke in the background of type 2 diabetes induces extensive remodeling of bone marrow communication networks, as characterized by reprogrammed signaling hierarchies and activation of pro-inflammatory ligand–receptor pathways.

### Diabetes and Stroke Amplify MIF, SIRP, and THBS Signaling within the Bone Marrow Niche

To further delineate changes in intercellular communication, we quantified ligand–receptor interaction strengths across conditions, focusing on HPC1 and monocytes based on the highlighted significant intercellular signaling from Figure 5B, which also have been previously identified as key signal-sending populations for hematopoiesis and myelopoiesis in diabetic stroke^44,45^. Bubble plots revealed significant MIF–(CD74+CXCR4), MIF–(CD74+CXCR2), and MIF–(CD74+CD44) interactions between HPC1 and other cell types, with markedly increased communication strength in diabetic (T2DM and T2DM.Stroke) groups relative to non-diabetic ones (Ctrl and Stroke) (Figure 6A). Diabetes specifically enhanced *Mif* and *CD74* expression in HPC1 and monocytes and granulocytes, respectively (Figure 6B and Supplementary Figure 3A). High constitutive expression of *Mif* and *CD74* were also found in HPC2 and HPC3 cells, respectively. Diabetes upregulated both *Mif* in HPC and *CD74* in monocytes/granulocytes regardless of stroke condition. Stroke consistently downregulated *CD74* expression in both diabetic and non-diabetic groups, although stroke displayed opposing effect on Mif in non-diabetic and diabetic mice (Figures 6C-D). Besides monocyte *CD74* gene expression, IF staining also confirmed the protein expression of CD74 on cell membrane of CD115+ monocytes in all groups, with representative images displayed in non-diabetic control mice (Figure 6E).

**Figure 6.**
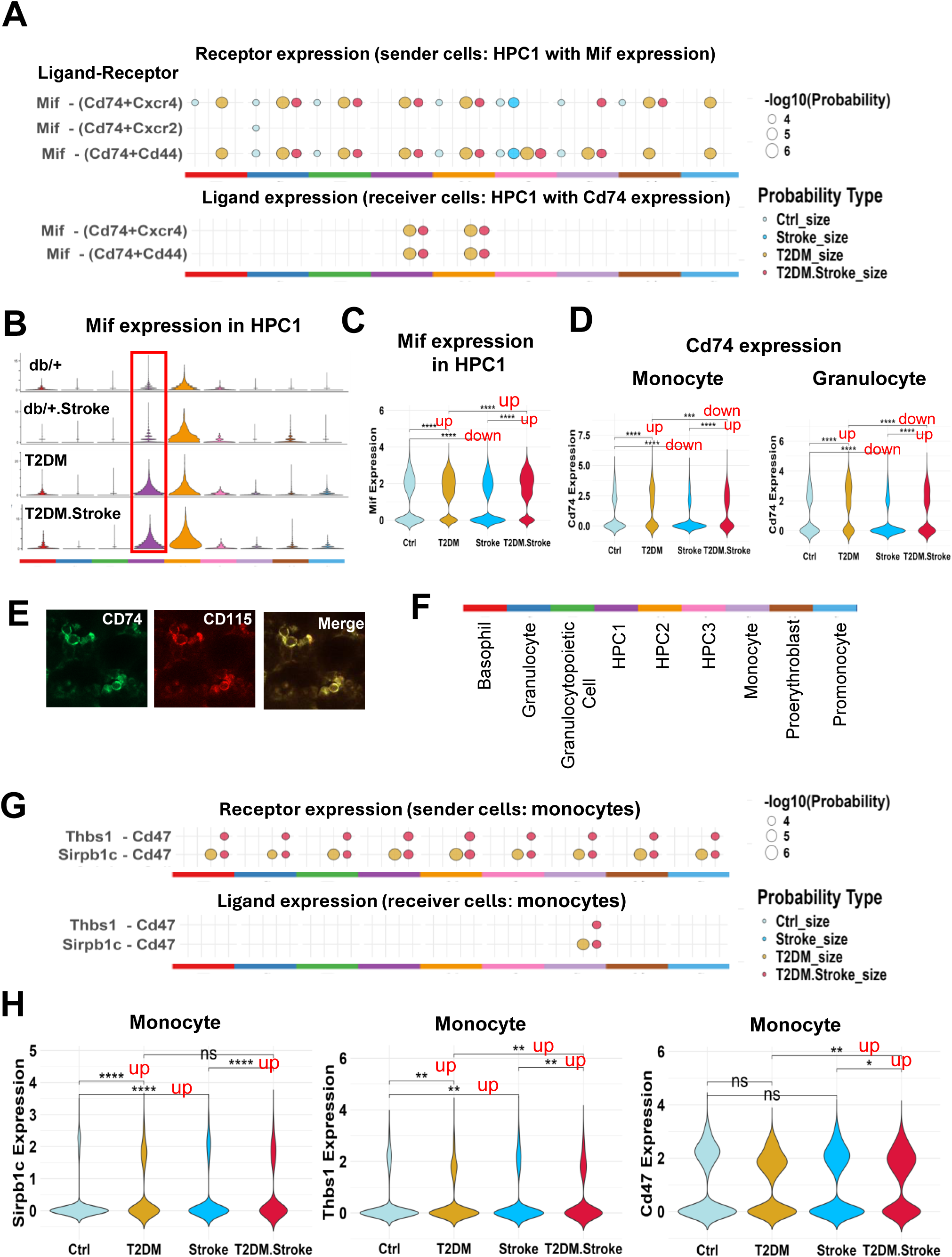
Altered ligand–receptor signaling and differential expression of MIF, SIRP, and THBS pathways in diabetic stroke bone marrow. **(A)** Bubble plots depicting significant MIF–CD74 ligand–receptor interactions between HPC1 cells and all other bone marrow cell clusters across the four experimental groups (Ctrl, T2DM, Stroke, and T2DM + Stroke). Colored circles represent different groups. Size of circles represents interaction probability of ligand-receptor pairs across groups. Stronger interactions are observed between HPC1 and other cell types in the T2DM and T2DM + Stroke groups. Color bar indicating each cell cluster shows in (**F**). **(B)** The distribution of Mif gene expression (ligand in MIF signaling) in all cell clusters across the four groups. Red boxes highlight the expression in HPC1 in the T2DM and T2DM + Stroke groups. **(C)** Violin plot shows statistic figure of Mif expression in HPC1. **(D)** Violin plot shows statistic figure of Cd74 expression in monocytes and granulocytes. **(E)** Representative immunofluorescence images showing CD74 expression in CD115+ monocytes in bone marrow of non-diabetic mice. **(F)** Color bar indicates each cell cluster. **(G)** Bubble plots showing SIRPB1C–CD47 and THBS1–CD47 ligand–receptor interactions between monocytes and other cell types. Monocytes expressing Sirpb1c and Thbs1 exhibit enhanced signaling interactions with other clusters in the T2DM and T2DM + Stroke groups. **(H)** Violin plots show statistic figures of Sirpb1c, Thbs1, and Cd47 expression in monocytes across conditions. Gene expression was analyzed using normalized, log-transformed data, with asterisks indicating statistical significance. For clearer visualization, significance was indicated by labeling as ‘up’ or ‘down’ for those showing significant increases or decreases, respectively.

Network visualization using CellChat circos plots (Supplementary Figure 4) further demonstrated intensified MIF signaling interactions among HPC1 and other hematopoietic clusters in diabetic groups (T2DM and T2DM.Stroke) (Supplementary Figure 4A), reflecting robust and disease-specific proinflammatory communication networks.

We next examined monocyte-derived SIRPB1C–CD47 signaling enriched in diabetic groups (T2DM and T2DM.Stroke) and THBS1–CD47 signaling enriched in diabetic mice after stroke (T2DM.Stroke) (Figure 6G). In diabetic groups (T2DM and T2DM.Stroke), monocytes expressing *Sirpb1c* and *Thbs1* exhibited enhanced outgoing signaling toward *Cd47*-expressing clusters. *Sirpb1c* and *Thbs1* were predominantly expressed in monocytes, whereas *Cd47* was broadly expressed across all bone marrow clusters (Supplementary Figures 3B–C). Violin plots confirmed significant upregulation of *Sirpb1c* and *Thbs1* in monocytes under diabetic conditions at both baseline and post-stroke, with *Cd47* expression also elevated in monocytes after diabetic stroke (Figure 6H). The SIRP pathway (Supplementary Figure 4B) showed dense outgoing interactions exclusively from monocytes in diabetic groups (T2DM and T2DM.Stroke), targeting all other cell populations. Similarly, the THBS1–CD47 signaling network (Supplementary Figure 4C) was markedly expanded in diabetic stroke marrow, with monocytes serving as the dominant source of signaling output.

Collectively, these findings support a model in which diabetes and stroke synergistically amplify intercellular communication within the bone marrow niche through MIF, SIRP, and THBS signaling axes, primarily driven by HPC1 and monocyte populations. These reprogrammed signaling interactions likely contribute to the inflammatory activation and functional remodeling of the diabetic bone marrow microenvironment following ischemic injury.

### Diabetes and Stroke Induce Metabolic Activation and Immune dysregulation in Hematopoietic and Myeloid Populations

To characterize transcriptional reprogramming of functional pathways in BM cells under diabetic and stroke conditions, we performed AUCell enrichment analysis to quantify gene set activity for metabolic and immune-related signatures across all experimental groups.

Glycolysis, adipogenesis, and oxidative phosphorylation pathways were significantly upregulated in diabetic groups (T2DM and T2DM.Stroke) relative to non-diabetic (Ctrl and Stroke) groups in HPC1, promonocyte/monocyte lineages (Figure 7A and C) as well as granulocytes and granulocytopoietic cells (Supplementary Figure 5A), indicating enhanced metabolic activation under diabetic conditions. Although previous studies found no significant change in rodent bone marrow adipocyte volume fraction, density or size after ischemic stroke^46,47^, our AUCell score shows stroke also increased a certain metabolic pathway activity in both genotypes, suggesting convergent metabolic stress responses to ischemic injury. These changes define a distinct metabolic profile associated with combined metabolic and ischemic stress. Diabetes downregulated IFN-γ signaling while stroke upregulated it only in the non-diabetic group in monocytes and promonocytes (Figure 7B and D, Supplementary Figure 6B). These findings suggest that diabetic stress attenuates immune responsiveness in HPC1, potentially altering differentiation or effector functions.

**Figure 7.**
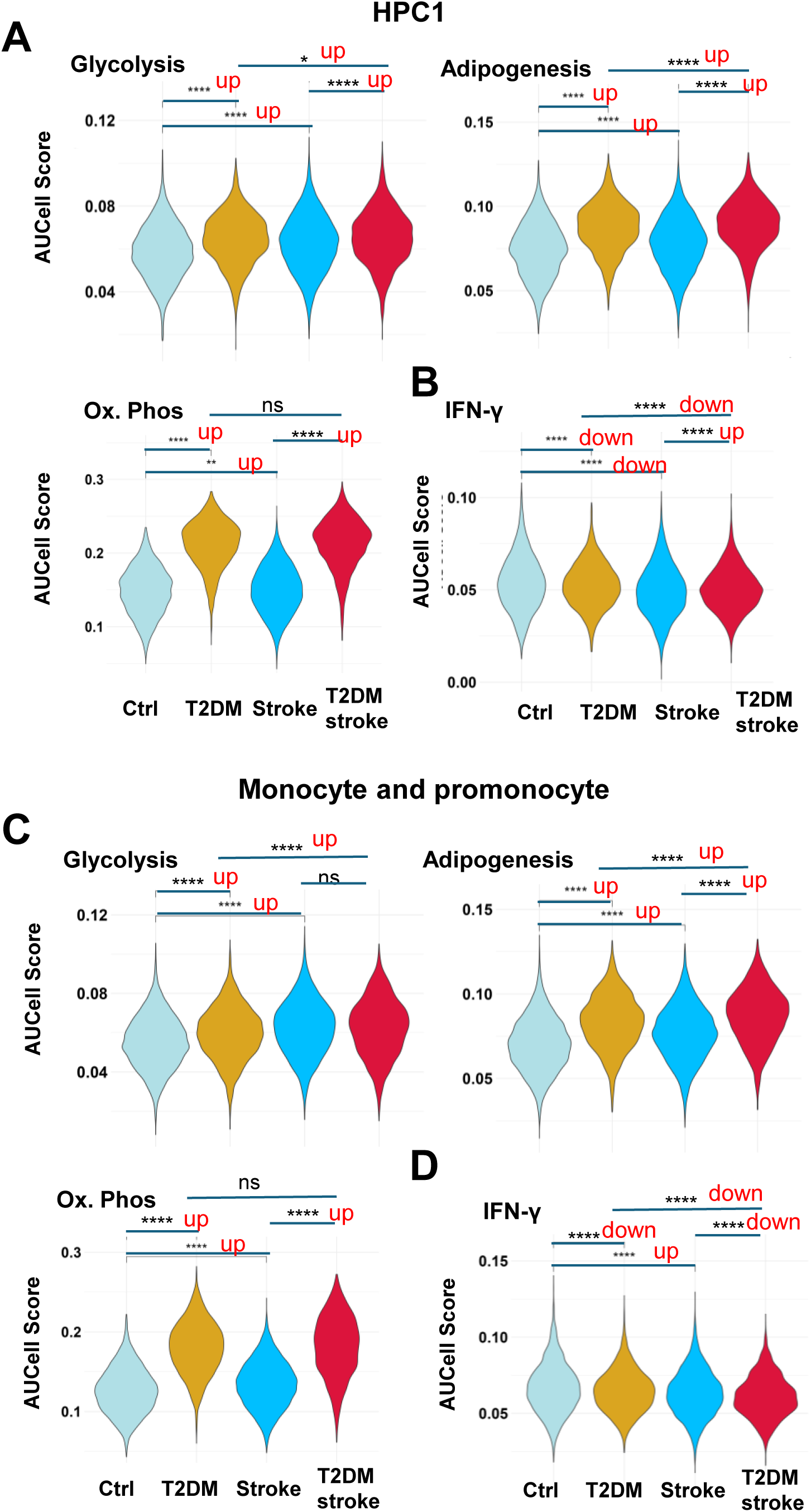
Metabolic and inflammatory pathway activity in HPC1 and monocyte/promonocyte populations across experimental groups. **(A)** AUCell-based signaling scores for metabolic pathway signatures (glycolysis, adipogenesis, and oxidative phosphorylation) in HPC1 cells from Ctrl, T2DM, Stroke, and T2DM + Stroke groups. **(B)** AUCell scores for IFN-Υ in HPC1 cells across the four groups. **(C)** Metabolic pathway signatures (glycolysis, adipogenesis, and oxidative phosphorylation) in monocyte/promonocyte populations across all experimental conditions. **(D)** IFN-Υ signature in monocyte/promonocyte populations across all experimental groups. AUCell scores were calculated as the area under the curve (AUC) for the ranked expression of pathway gene sets in each individual cell. 10,000 down-sampled whole BM cells, 15,000 HPC1 and 17,609 monocytes/promonocytes across all four experimental groups were used for AUCell analysis. Statistical comparisons between groups were performed using Wilcox test, with asterisks indicating statistical significance.

Together, these results demonstrate a coordinated transcriptional shift toward metabolic activation and immune dysregulation in BM cells under diabetic stroke conditions. This reprogramming reflects an altered bone marrow microenvironment that promotes myeloid lineage bias and inflammatory activation while compromising specific immune responsiveness following ischemic injury.

### Diabetic Stroke Enhances Myeloid Activation and Migratory Programs in Bone Marrow CD115^⁺^ Monocytes

To assess the effects of diabetes and stroke on bone marrow immune cell composition and activation, we performed GeoMx digital spatial profiling (DSP) to spatially capture and quantify gene expression in monocyte (CD115⁺) and neutrophil (Ly6G⁺) populations across experimental groups (Figure 8A).

Differential gene expression (DEG) analysis of CD115⁺ monocytes and Ly6G⁺ neutrophils revealed distinct transcriptional responses under diabetic and stroke conditions. Scatter plots comparing log fold changes between db/db stroke vs db/db (x-axis) and db/+ stroke vs db/+ (y-axis) identified subsets of genes that were consistently upregulated by stroke in both genotypes (red dots) as well as genes uniquely elevated only in the diabetic groups comparison (db/db stroke vs db/db, green dots) (Figure 8B). These distributions indicate both shared and diabetes-specific transcriptional adaptations in myeloid cells.

Gene Ontology (GO) enrichment analysis of shared upregulated genes by stroke in both genotypes highlighted biological processes (BP) related to neutrophil activation, myeloid leukocyte activation, and injury response, including antimicrobial humoral defense, response to bacterial and toxic stimuli, and regulation of apoptotic and immune effector mechanisms (Figure 8C). Biological processes uniquely upregulated by stroke in the diabetic mice (db/db stroke vs db/db) revealed enrichment of translational and migratory programs such as ribosome assembly, cytoplasmic translation, leukocyte cell–cell adhesion, and leukocyte and myeloid migration (Figure 8D).

**Figure 8.**
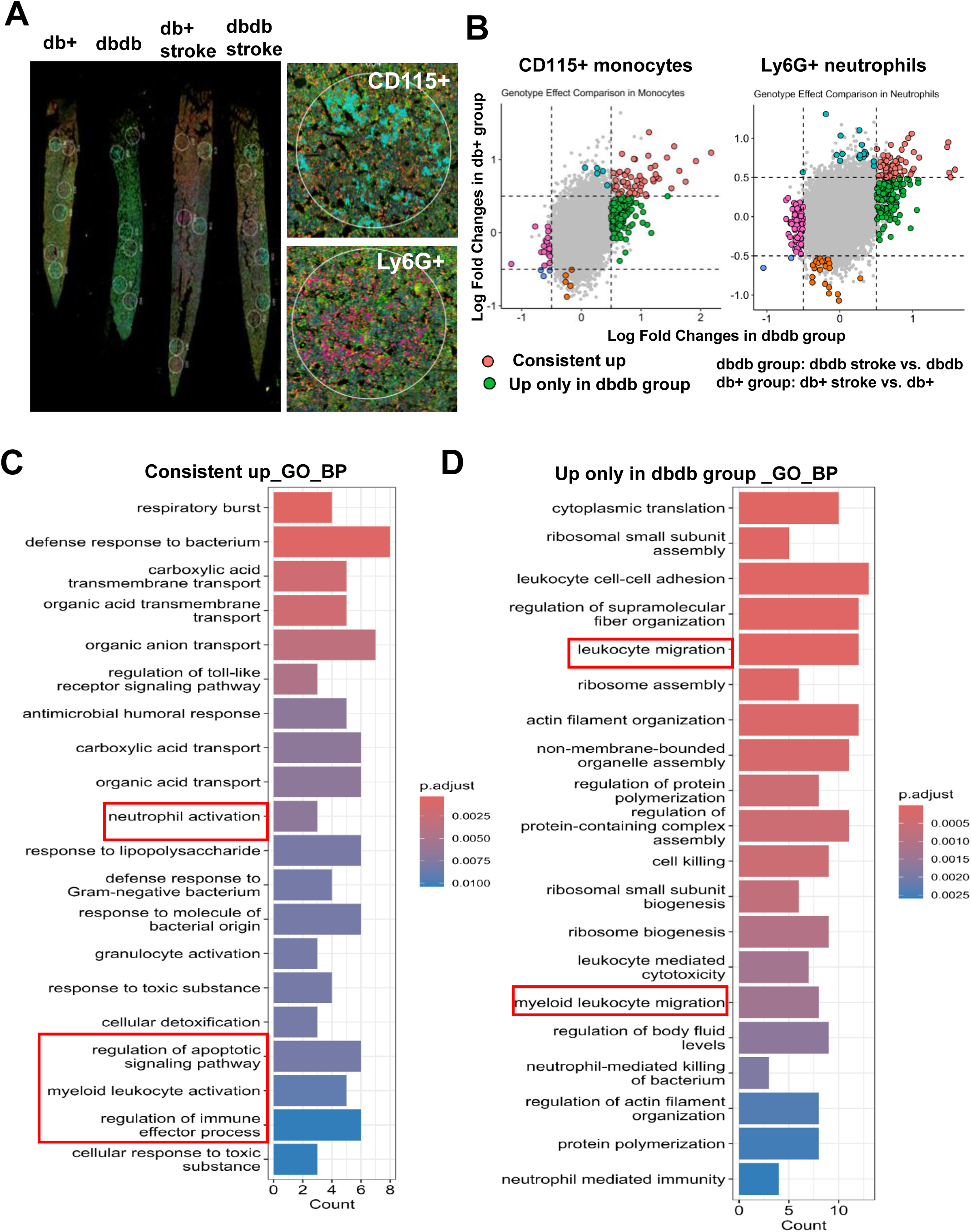
Spatial profiling of CD115^⁺^ monocytes and Ly6G^⁺^ neutrophils in control and diabetic mice with or without stroke. **(A)** Representative immunofluorescence images of CD115 and Ly6G staining in whole bone marrow sections from the four experimental groups (db/+, db/db, db/+ stroke, and db/db stroke), along with examples of cell type segmentation with randomly selection within bone marrow section. **(B)** Volcano plots showing differentially expressed genes (DEGs) in CD115⁺ monocytes and Ly6G⁺ neutrophils between stroke and non-stroke conditions in each genotype. The X-axis represents log₂ fold change in dbdb stroke vs. dbdb (dbdb group), and the Y-axis represents db+ stroke vs. db+ (db+ group). Fisher’s exact test was used for the analysis, with cutoff thresholds of FDR = 0.05. Dashed lines at ±0.5 log₂ fold change indicates the threshold for significant gene expression changes. Each dot represents a gene. Gray dots denote non-significant genes. Red dots indicate shared genes that significantly upregulated both group comparisons (db+ group and dbdb group). Green dots indicate genes significantly upregulated in dbdb group comparison. **(C)** GO biological process (BP) enrichment analysis of DEGs consistently upregulated in CD115⁺ monocytes in both db/+ and db/db group comparisons. **(D)** GO BP enrichment analysis of DEGs uniquely upregulated in CD115⁺ monocytes from the db/db comparison group.

These findings suggest that diabetes-specific cues drive a transcriptional program in monocytes characterized by enhanced protein synthesis and migratory potential, potentially predisposing them to extramedullary trafficking or heightened inflammatory activation.

### Diabetes and Stroke Synergistically Amplify Inflammatory Programs in Bone Marrow Neutrophils

Similar approach was applied to the analysis of neutrophils. Genes consistently upregulated in Ly6G⁺ neutrophils across diabetic and stroke conditions (db/db stroke vs db/db on the x-axis, db/+ stroke vs db/+ on the y-axis) showed strong enrichment for innate immune activation, myeloid cell migration, activation, and chemotaxis (Supplementary Figure 6A), reflecting a shared inflammatory program under both pathological contexts. Notably, GO terms uniquely enriched in the diabetic comparison (db/db stroke vs db/db) included leukocyte degranulation, generation of precursor metabolites and energy, and calcium ion transport (Supplementary Figure 6B), suggesting that diabetes specifically primes neutrophils for enhanced degranulation and metabolic adaptation, a response absent in stroke alone.

### Diabetes and Stroke Amplify Myeloid Inflammatory Gene Programs in Bone Marrow

To validate these pathway-level findings, we examined highly shared genes that enriched in GO terms of leukocyte activation and migration using GeoMx DSP data from sorted CD115⁺ monocytes and Ly6G⁺ neutrophils (Figure 9A). Diabetes significantly elevated the expressions of *Elane, Mpo, S100a9, Camp, Cd47, Cd177 and Anxa1*relative to non-diabetic ones, at either baseline or after stroke. These genes are integral to GO pathways associated with myeloid activation, migration, and chemotaxis.

**Figure 9.**
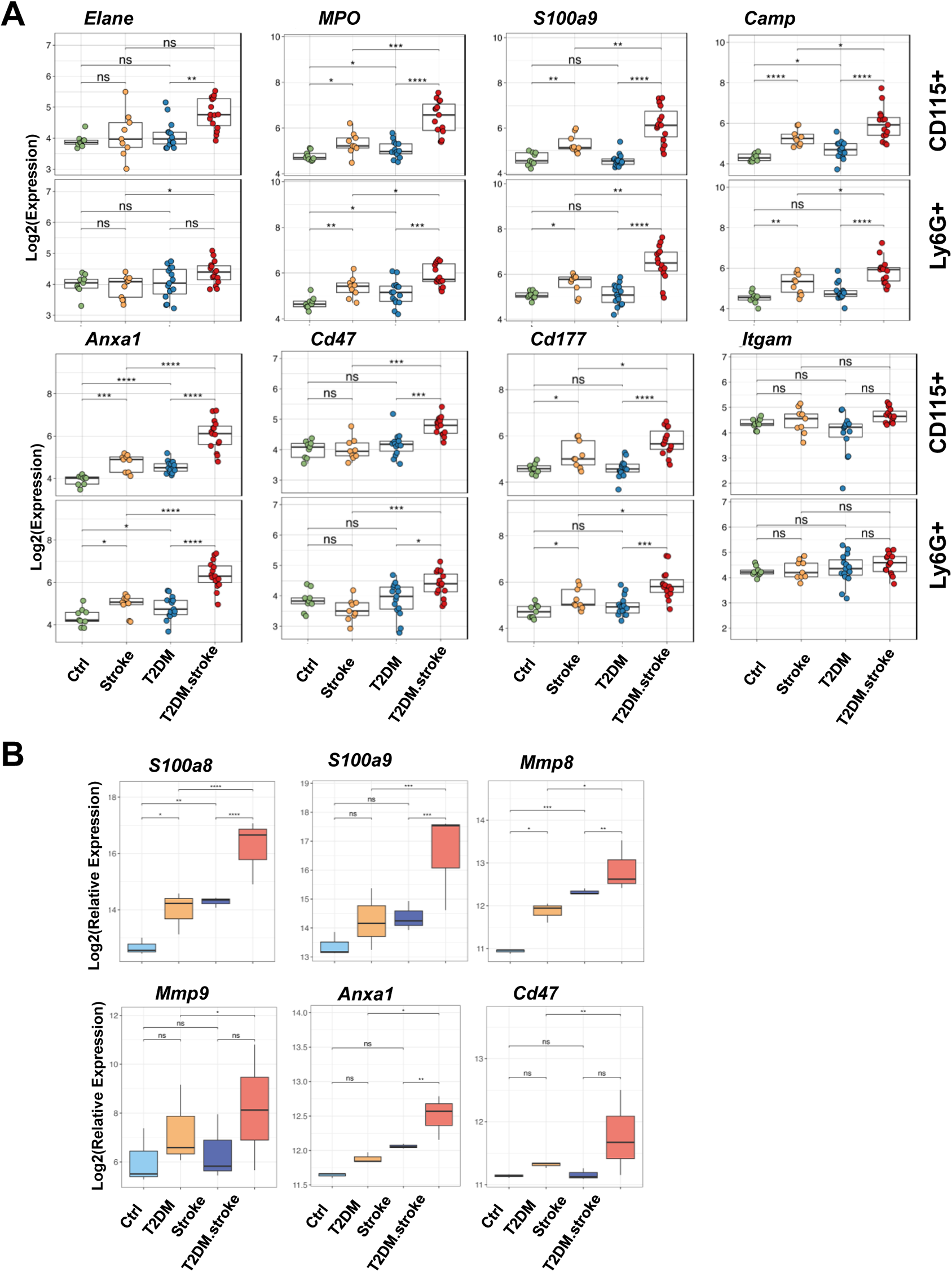
Proinflammatory factor gene expression from GeoMx DSP and nCounter. **(A)** Gene expression of leukocyte activation and migration–related genes in CD115+ monocytes and Ly6G+ neutrophils from selected ROIs in bone marrow, measured by GeoMx Digital Spatial Profiling (DSP). Genes include *Elane, Mpo, S100a9, Camp, Anxa1, Cd47, Cd177,* and *Itgam*. Data is shown for four experimental groups: db+, dbdb, db+ stroke, and dbdb stroke. Fisher’s exact test was used for the analysis, with cutoff thresholds of FDR = 0.05. Asterisks indicate statistical significance. **(B)** Gene expression in bone marrow monocytes measured by nCounter for *S100a8, S100a9, Mmp8, Mmp9, Anxa1 and Cd47* across the same four experimental groups. The gene expression is normalized using the geometric mean of housekeeping genes to account for technical variability across regions of interest and groups. Fisher’s exact test was used for the analysis, with cutoff thresholds of FDR = 0.05. Asterisks indicate statistical significance.

We further validated gene expression in monocytes isolated from bone marrow using nCounter (Figure 9B). Pro-inflammatory and effector genes *S100a8, S100a9, Mmp8, Mmp9, Anxa1, and Cd47*, key mediators of neutrophil and monocyte activation, were significantly upregulated by stroke in diabetic mice. *S100a8, Mmp8 were also increased by diabetes in non-stroke groups, or by stroke in non-diabetic groups. S100a8, S100a9, Mmp8 and Anxa1* were markedly elevated by diabetes in stroke groups.

Taken together, these results demonstrate that diabetes and stroke synergistically reprogram bone marrow myeloid cells, enhancing inflammatory signaling, matrix remodeling, and immune effector gene expression. Diabetes uniquely augments transcriptional programs related to neutrophil degranulation and leukocyte migration, and these effects are further amplified by ischemic stress. This diabetes-specific priming of the bone marrow myeloid compartment likely contributes to the sustained inflammatory activation and immune dysregulation characteristic of diabetic stroke.

## DISCUSSION

### Bone marrow atrophy and lineage bias in diabetic stroke

The current study identifies BM as a critical regulator of immune dysfunction in diabetic stroke and defines how chronic metabolic disease exacerbates post-stroke pathology. By integrating single-cell transcriptomics, digital spatial profiling, and functional validation, we demonstrate that T2DM remodeled the BM microenvironment, disrupted hematopoietic homeostasis, and primed myeloid lineages for maladaptive responses to ischemic injury. We found that T2DM induced significant BM atrophy and cellular depletion, accompanied by a shift in hematopoietic lineage commitment. Early lymphoid progenitors were markedly reduced, and lymphoid differentiation was suppressed, consistent with prior studies linking chronic hyperglycemia to HSC exhaustion and niche dysfunction^48–50^. These alterations create a dysfunctional marrow environment incapable of sustaining balanced immune output ^51^. Although ischemic stroke alone produced mild architectural disruption, stroke and diabetes synergistically reduced BM cellularity and dysregulated hematopoiesis, indicating that diabetic BM is primed to amplify injury-induced dysfunction^52,53^.

### Disrupted myeloid development and uncoupling of differentiation and maturation

Trajectory analyses revealed a disordered myeloid development in the diabetic BM, which was further exacerbated by stroke. Diabetes drives a shift toward later pseudotime accompanied by increased transcriptional heterogeneity across myeloid lineages, suggesting metabolic stress primes hematopoietic cells toward a more developed or activated transcriptional state even in the absence of acute injury, potentially reflecting chronic low-grade inflammation^54,55^.

Differentiation and maturation signatures become uncoupled, most prominently in whole BM myeloid cells, HPC1 and granulocytes in diabetes and in monocytes in diabetic after stroke. Stroke in non-diabetes triggered coordinated changes in hematopoietic trajectories, marked by increased differentiation and maturation, which is consistent with emergency myelopoiesis^6^, in which the bone marrow rapidly increases myeloid production in response to acute injury^56,57^. In contrast, diabetes markedly disrupted this adaptive response. Although diabetic bone marrow cells were shifted toward later pseudotime values and displayed increased heterogeneity, T2DM animals exhibited reduced differentiation potential at baseline in progenitor compartments, suggesting impaired early lineage commitment despite apparent transcriptional advancement. When stroke occurred in the diabetic setting, the expected stroke-induced amplification of differentiation and maturation programs was blunted or lost across bone marrow, HPC1, and granulocytic populations. This uncoupling of developmental progression from functional maturation indicates that diabetes compromises the bone marrow’s capacity to mount a coordinated and effective myeloid response following ischemic injury. Notably, monocytes represented an exception to this global impairment. In diabetes, monocytes displayed increased differentiation and maturation signatures at baseline and maturation following stroke, consistent with a more activated or developmentally advanced state. This favoring monocyte activation in chronic metabolic inflammation, contributing to maladaptive inflammatory responses in downstream compartments, including peripheral blood and ischemic brain tissue^58^. Together, these data support a model in which diabetes imposes a chronic reprogramming of bone marrow hematopoiesis that alters developmental timing, increases cellular heterogeneity, and reduces functional plasticity.

### Reprogrammed bone marrow communication networks in diabetes and stroke

Cell–cell communication analysis uncovered a shift in signaling hierarchies, with HPC1 and monocytes emerging as dominant hubs enriched for MIF, SIRP, and THBS signaling. These ligand–receptor networks are implicated in immune evasion, progenitor stress, and sterile inflammation, suggesting a transition from regenerative to inflammatory intercellular communication within the diabetic BM niche^59,60^. Importantly, the upregulation of these signaling pathways after ischemic stroke has been linked to worsened stroke outcomes^54,61–63^. Although previous studies have shown that MIF promotes neurological recovery after ischemic stroke^64,65^, excess MIF expression has been linked to exaggerated inflammation and immunopathology^66^. MIF–CD74 signaling enhances monocyte survival and promotes sustained NF-κB activation, leading to increased production of pro-inflammatory cytokines and resistance to anti-inflammatory cues^67^. In the post-stroke setting, this results in exaggerated monocyte-mediated inflammation, disruption of the BBB, and heightened neuronal injury^62^. The SIRP–CD47 pathway, typically functioning as a ‘don’t eat me’ signal, becomes dysregulated, suppressing clearance of activated or damaged cells and promoting immune evasion. This hampers resolution of inflammation and permits the persistence of dysfunctional immune cells^60^. Furthermore, THBS1–CD47 signaling impairs nitric oxide signaling and endothelial function, contributing to vascular dysfunction, leukocyte adhesion, and reduced cerebral perfusion^68^. Together, these interactions exacerbate neuroinflammation, compromise tissue repair, and increase the risk of secondary injury in diabetic stroke^54,55^.

### Metabolic activation with immune dysregulation in progenitors and myeloid cells

Transcriptional profiling further demonstrated a decoupling of metabolic and immune responses in hematopoietic and myeloid populations. These findings support a model of metabolic priming, in which diabetic progenitor and myeloid cells exhibit elevated energy production but diminished immune responsiveness^22,69^. The observed reduction in inflammatory signatures in diabetes is consistent with features of immune exhaustion^70^, which arises in the context of chronic inflammatory or antigenic stimulation^71^. In this state, inflammatory genes may no longer rank among the most highly expressed within each individual cell calculated by AUCell function, resulting in reduced inflammatory signature scores even as baseline inflammatory signaling persists at the tissue level^72,73^.

### Diabetes specific priming of myeloid effector programs

Spatial transcriptomic analysis of Ly6G⁺ neutrophils and CD115⁺ monocytes and nCounter analysis of isolated monocytes from BM provided additional evidence of diabetes-specific transcriptional priming. These cells exhibited increased expression of *MPO, S100a8*, *S100a9*, *Mmp8*, *Mmp9*, *Anxa1* and Cd47 in diabetic groups, which involved in enhanced degranulation, chemotaxis, and matrix remodeling programs. These molecular patterns parallel findings from peripheral immune profiling of diabetic stroke patients, who display elevated circulating neutrophil and monocyte activation markers and heightened oxidative and proteolytic activity^74^. The enhanced migratory and degranulation signatures in the diabetic BM suggest that these cells are preconditioned for hyperreactive responses upon mobilization, promoting vascular inflammation and secondary injury after stroke. Several studies report context-dependent roles for macrophage migration inhibitory factor (MIF) in ischemic injury^75^. MIF signaling protects against ischemia/reperfusion damage by enhancing AMPK and survival pathways in cardiomyocytes^76^ or reducing oxidative stress and caspase-3 activation in neurons^77,78^. Conversely, in other contexts MIF contributes to pathogenic inflammation and leukocyte recruitment, and deletion or inhibition of MIF reduces infarct size and neurologic deficits in experimental stroke, suggesting that excessive MIF can amplify inflammatory cell infiltration and oxidative damage^79^. We also observed Mif expression trended higher in diabetes, although no significance (data not shown), supporting MIF as a candidate mediator of dysregulated myeloid priming in diabetes. Together, the BM signatures support a primed myeloid state in diabetes that may heighten inflammatory and proteolytic responses after mobilization, a hypothesis that requires direct testing in blood and brain compartments.

### Implications for diabetic stroke pathology and therapy

Collectively, these findings support a model in which diabetes establishes a maladaptive BM niche characterized by metabolic stress, aberrant signaling, and immune imbalance. This preconditioned environment limits adaptive the hematopoietic response to ischemic injury, amplifying inflammatory cascades and compromises impairing tissue repair after stroke.

From a translational perspective, the diabetic bone marrow emerges as an upstream regulator of post stroke inflammation and a viable therapeutic target. Interventions aimed at restoring hematopoietic balance, normalizing marrow metabolism, or reprogramming myeloid differentiation could mitigate systemic inflammation and improve stroke recovery in diabetic patients. Prior preclinical studies report improved outcomes after ischemia reperfusion or stroke when targeting MIF or CD47, for example with ISO-1 (MIF inhibitor) or CD47 blocking antibodies. Our findings nominate these pathways for mechanistic testing in diabetic stroke settings^64,68,80^. In parallel, metabolic reprogramming of hematopoietic stem/progenitor cells through PPAR-δ or PPAR-γ signaling^81,82^ has been shown to counteract maladaptive programming induced by chronic hyperglycemia.

Together, we define a direct mechanistic link between T2DM-derived BM dysfunction and worsened ischemic outcomes, positioning hematopoietic remodeling as a key determinant of diabetic stroke pathology and recovery.

## Acknowledgements

This work was supported by American heart Association 24POST1187683 (HZ), NIH 1R01NS137082-01A1 (JL), VA Merit Award 2I01BX003335 (JL), Research Career Scientist award 2IK6BX004600 (JL).

## SUPPLEMENTARY FIGURES

**Supplementary Figure 1.**
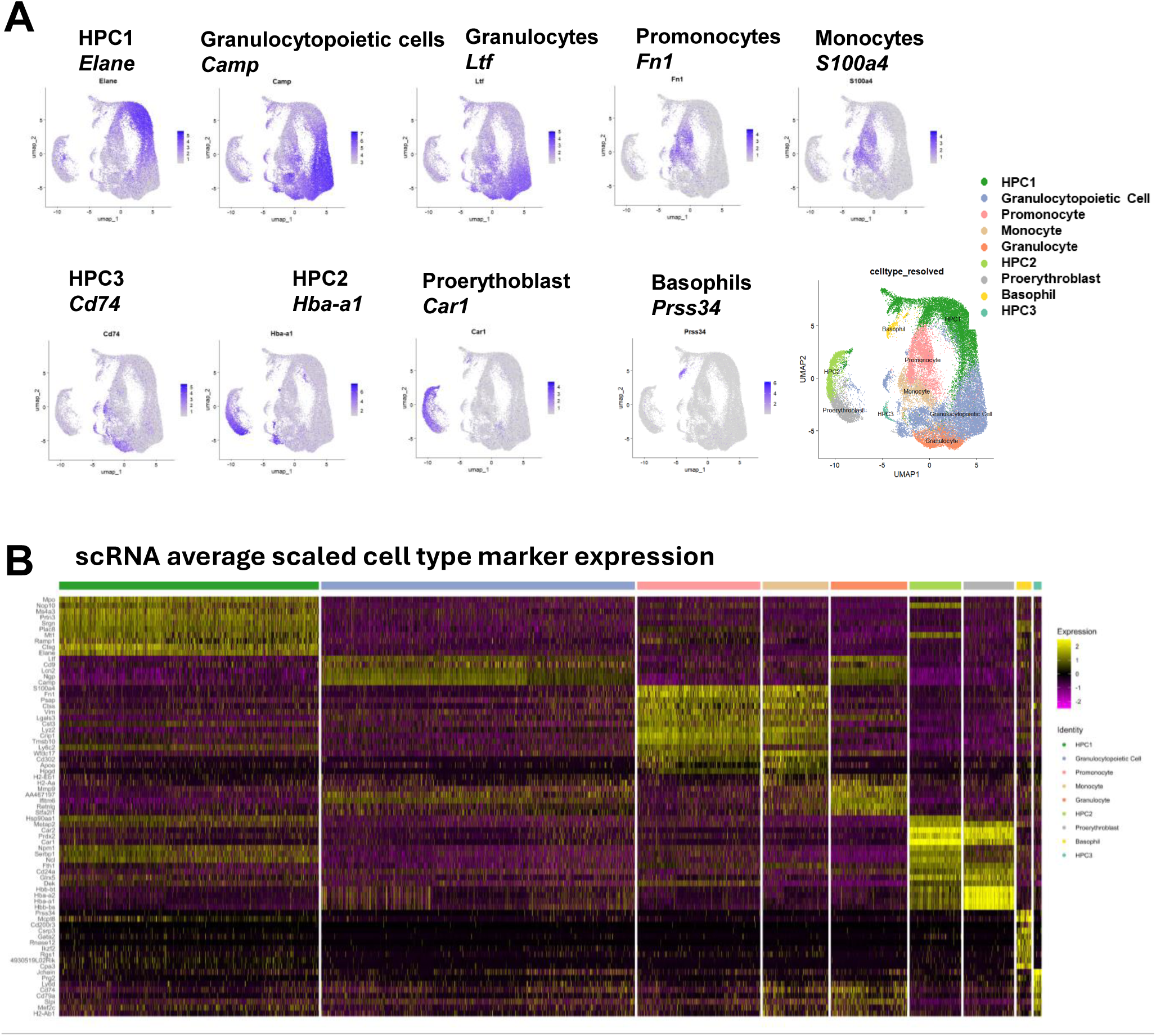
Gene markers identified for cell cluster annotation in bone marrow sc RNA sequencing. **(A)** Feature plots of representative gene marker of each cell cluster in UMAP. **(B)** Heatmap showing normalized gene expression (row-scaled per gene) for top and/or selected differentially expressed marker genes across annotated bone marrow cell types. The expression patterns reveal robust lineage-specific transcriptional signatures.

**Supplementary Figure 2.**
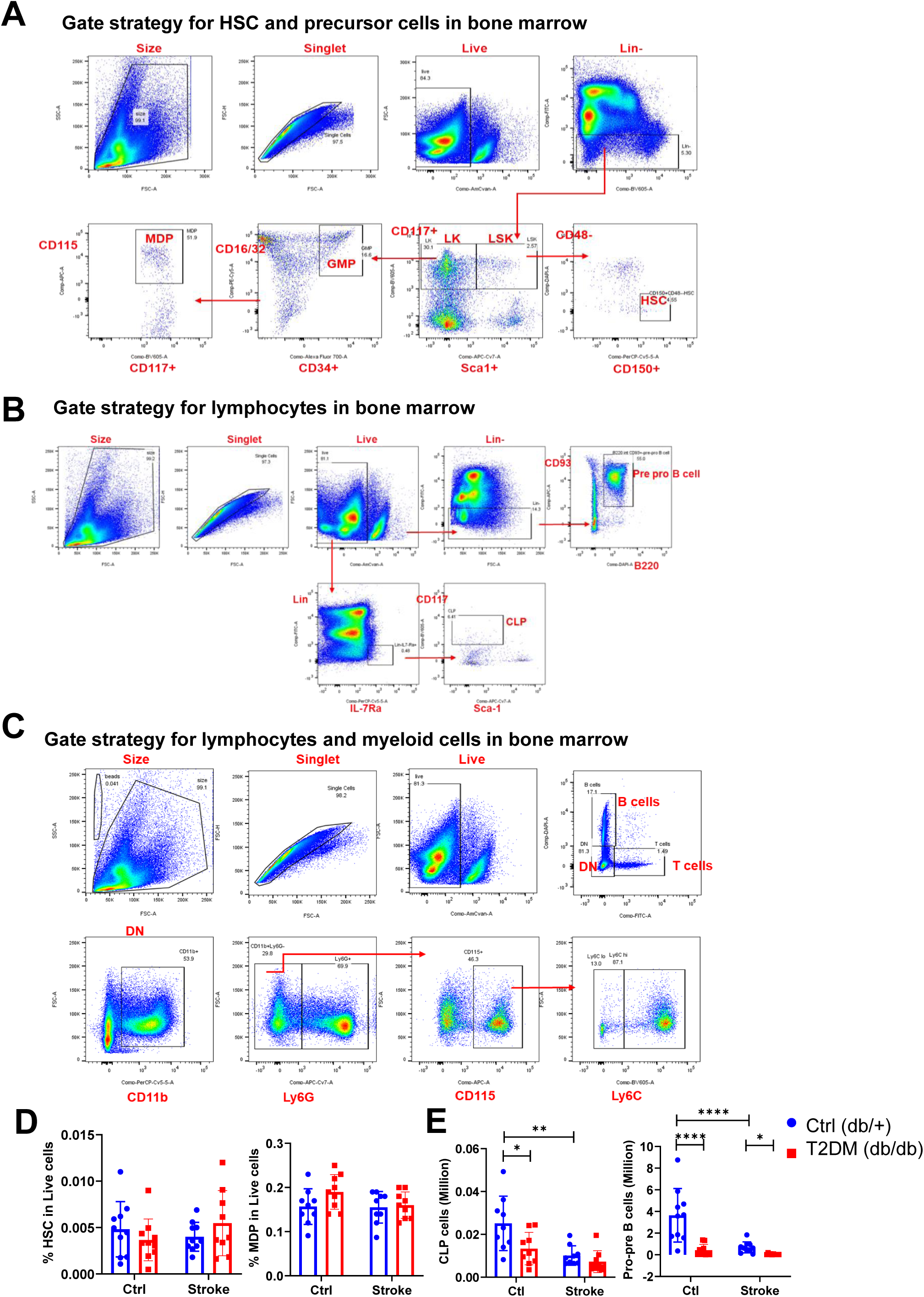
Gating strategy for FACS analysis of bone marrow populations. **(A)** Gating strategy for identifying hematopoietic stem cells (HSCs) and precursor populations. **(B)** Gating strategy for lymphocyte populations. **(C)** Gating strategy for myeloid populations. **(D)** Proportional changes in HSCs and monocyte–dendritic cell progenitors (MDPs) in live cells across four experimental groups (db/+, db/db, db/+ stroke, and db/db stroke). **(D)** Cell counts of CLP and pre-pro B cells in the bone marrow across the four groups. N= 8-10. Two-way ANOVA followed by Tukey’s post hoc test was used for analysis. Statistical significance is indicated as follows: \**P* < 0.05; \*\**P* < 0.01; \*\*\**P* < 0.001, \*\*\*\**P* < 0.0001.

**Supplementary Figure 3.**
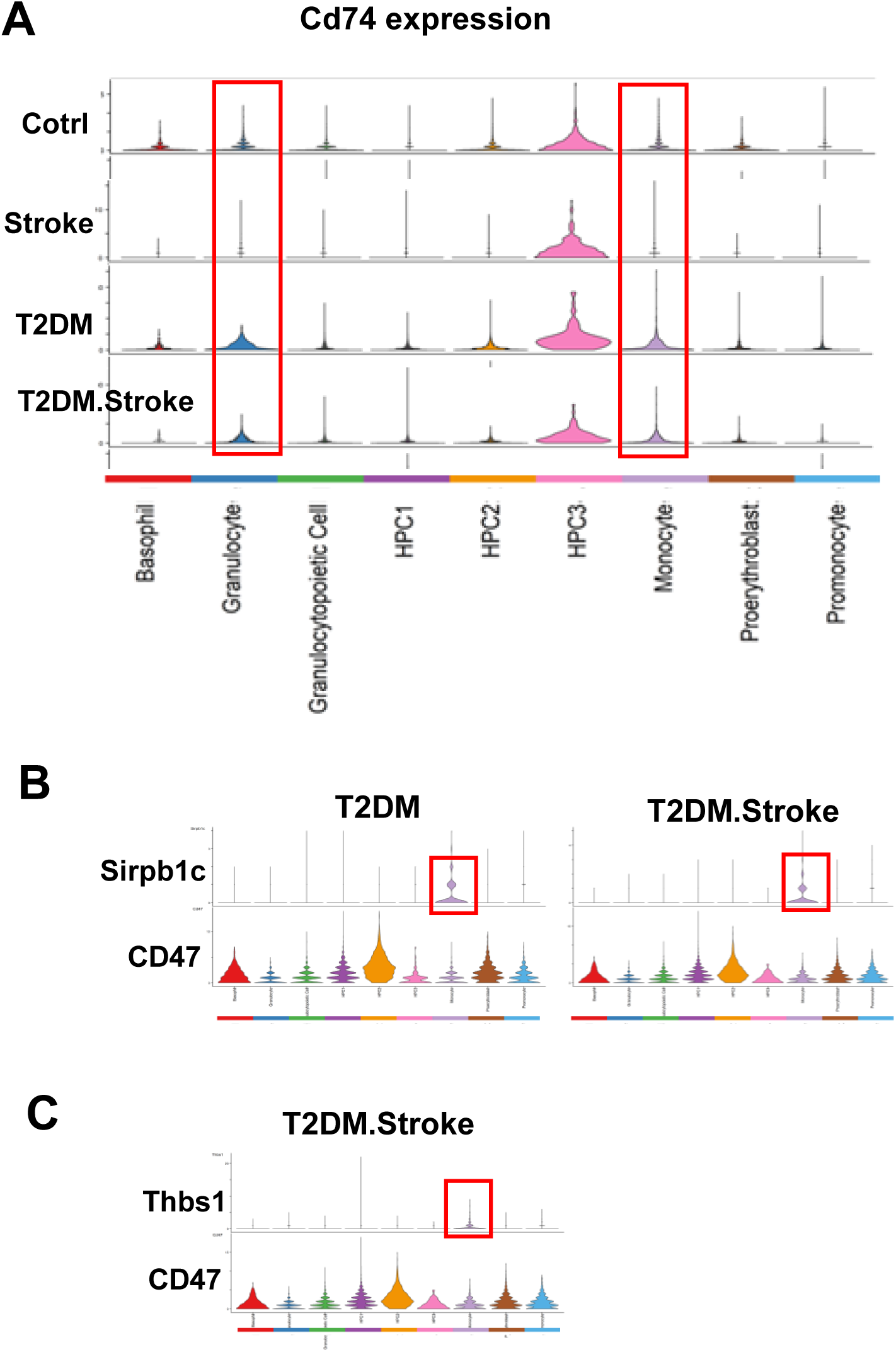
Expression patterns of MIF, SIRP, and THBS signaling pathway components in the bone marrow. **(A)** The distribution of Cd74 expression (a MIF receptor) in all cell clusters across all four experimental groups. Red boxes highlight expression in granulocytes and monocytes in T2DM and T2DM + *Stroke* groups. **(B)** The distribution of Sirpb1c and Cd47 expressions in all cell clusters across all four experimental groups. Red boxes highlight expression of ligands Sirpb1c in monocytes in T2DM and T2DM + Stroke groups. **(C)** The distribution of Thbs1 and Cd47 expression in all cell clusters across all four experimental groups. Red boxes highlight expression of ligands Thbs1 in monocytes in T2DM + Stroke groups.. Only specific groups showing statistically significant interactions identified by CellChat are displayed.

**Supplementary Figure 4.**
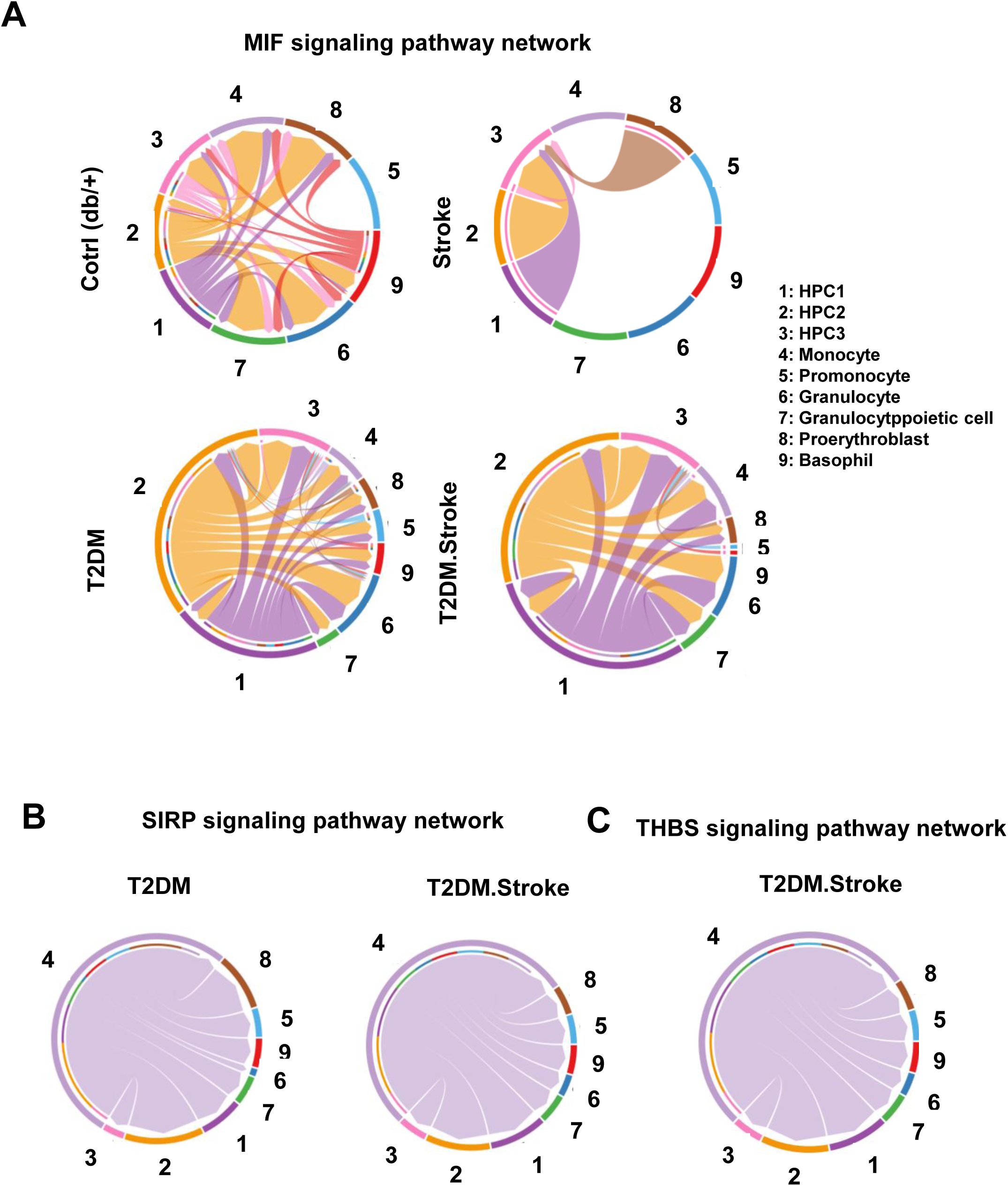
Altered intercellular communication networks involving HPC1 and monocytes across conditions. **(A)** Chord diagrams representing inferred MIF signaling interactions between HPC1 and other bone marrow cell populations in Ctrl, Stroke, T2DM, and T2DM + Stroke groups. **(B)** Chord diagrams showing SIRP and THBS signaling networks centered on monocytes in the T2DM and T2DM + Stroke groups. Number with colored node represents individual cell types, while directed chords indicate ligand–receptor–mediated signaling from source (sender) to target (receiver) populations. Chord thickness reflects the relative communication probability, and colors denote the signaling source population. Only statistically significant interactions identified by CellChat are displayed.

**Supplementary Figure 5.**
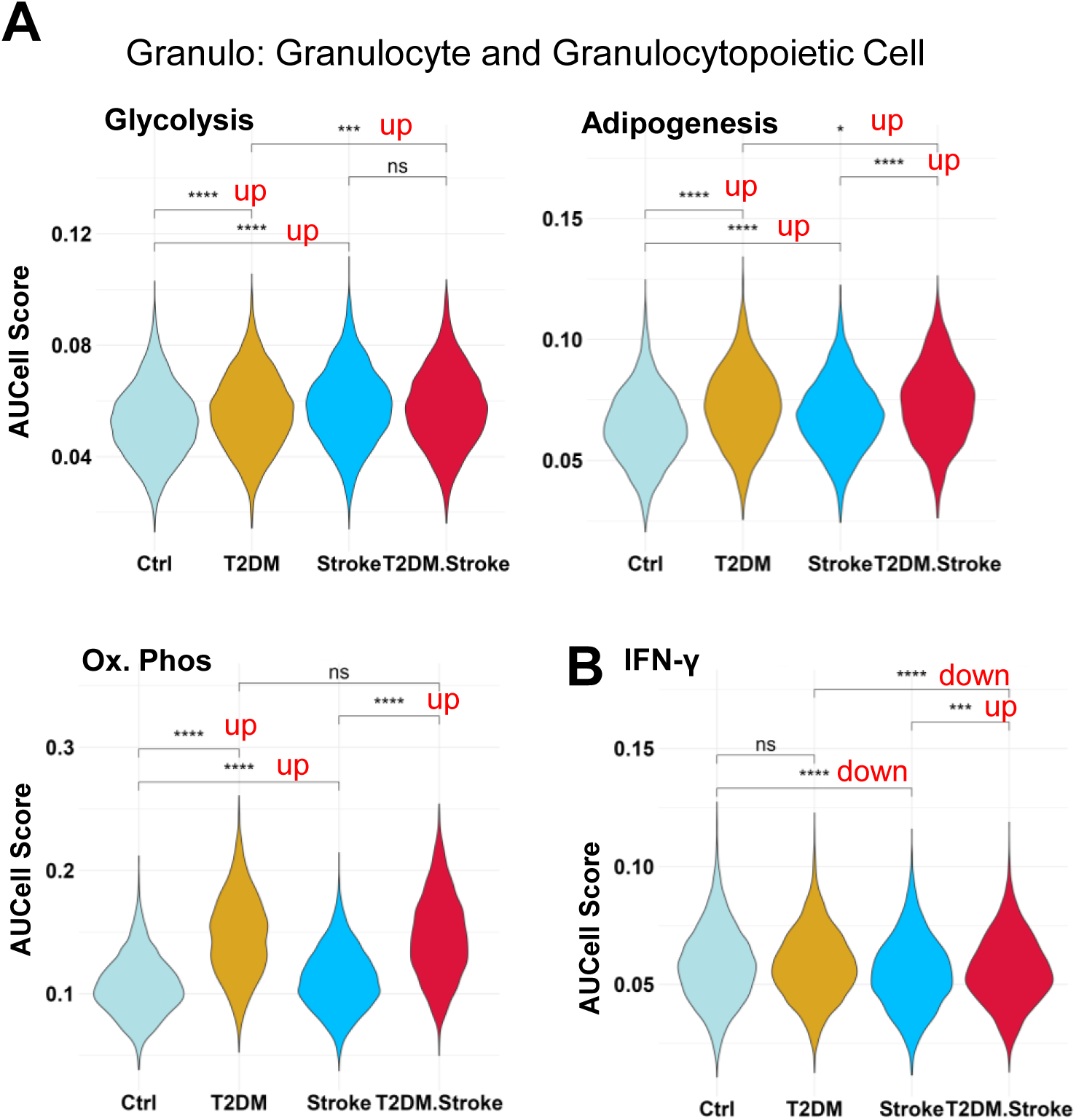
Metabolic and IFN-Υ signatures in granulocyte and granulocytopoietic cells. **(A)** AUCell scores for metabolic pathway signatures (glycolysis, adipogenesis, and oxidative phosphorylation) in granulocyte and granulocytopoietic cells across four groups: Ctrl, T2DM, Stroke, and T2DM + Stroke. **(B)** AUCell scores for IFN-Υ in granulocyte and granulocytopoietic cells across the four groups. AUCell scores were calculated as the area under the curve (AUC) for the ranked expression of pathway gene sets in each individual cell. 13,121 granulocytes/granulocytopoietic cells across all four experimental groups were used for AUCell analysis. Statistical comparisons between groups were performed using Wilcox test, with asterisks indicating statistical significance.

**Supplementary Figure 6.**
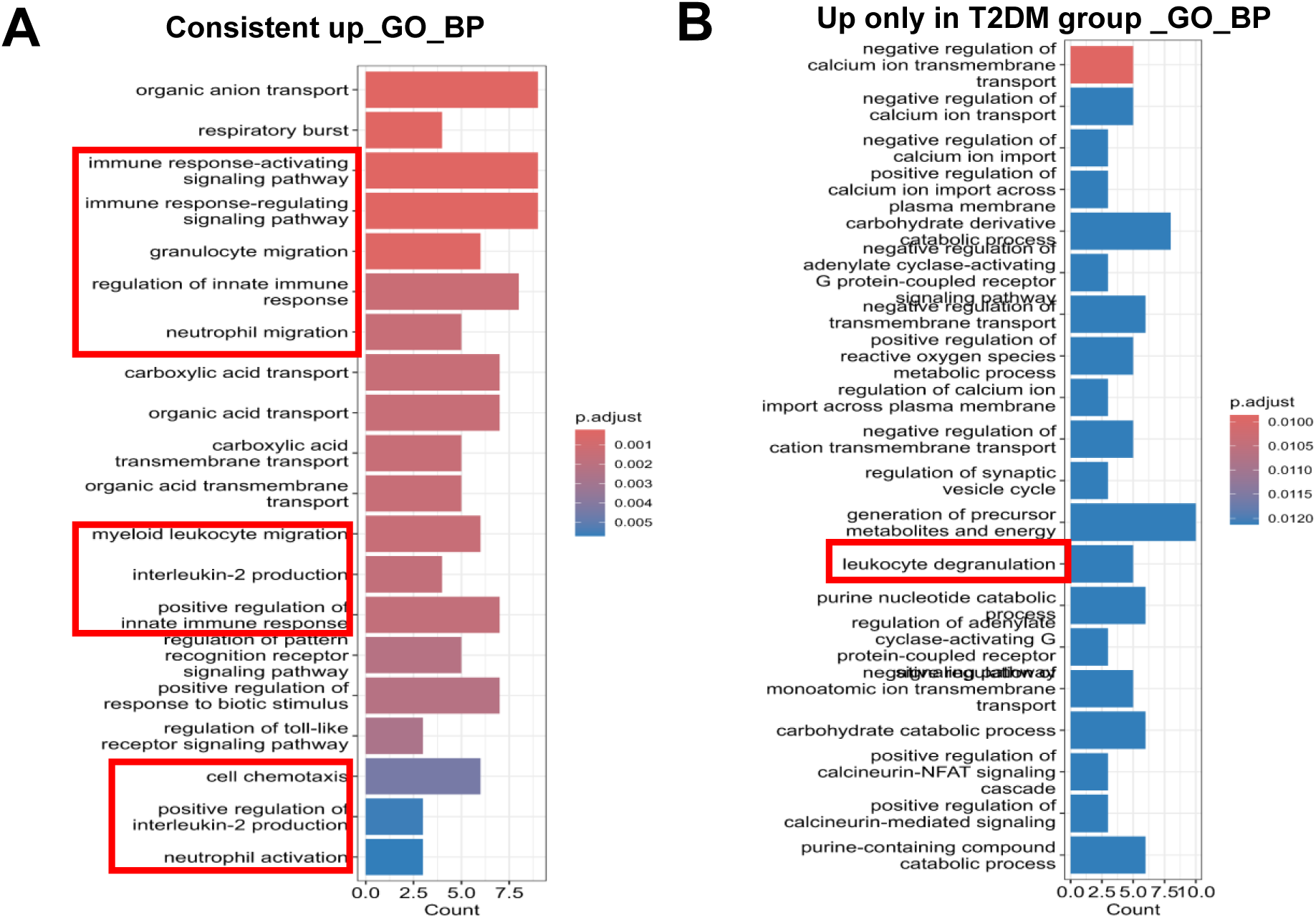
GeoMx DSP profile of GO Enrichment pathway analysis involved in biological process (BP) in Ly6G+ neutrophils. **(A)** Bar plots show the GO Enrichment pathway analysis involved in biological process (BP) of consistent up DEGs (up regulated both in dbdb group and db+ group) in Ly6G+ neutrophils. **(B)** Bar plots show the GO Enrichment pathway analysis involved in biological process (BP) of DEGs that are only upregulated in dbdb group in Ly6G+ neutrophils.

