## Supplemental Table 1 for "Type 2 diabetes Reprograms Bone Marrow Hematopoiesis and Dysregulates Immune Signaling in Response to Stroke"

**SingleR**

**Seurat Clusters**

|  | **Basophil** | **Granulocyte** | **Granulocytopoietic** | **HPC1** | **HPC2** | **HPC3** | **Monocyte** | **Proerythroblast** | **Promonocyte** |
| --- | --- | --- | --- | --- | --- | --- | --- | --- | --- |
| **B** | 0.62% | 5.76% | 1.92% | 0.77% | 2.66% | 25.58% | 1.36% | 2.33% | 0.39% |
| **Basophil** | 71.61% | 0.25% | 0.18% | 1.28% | 0.05% | 0.27% | 0.11% | 0.19% | 0.00% |
| **Erythroblast** | 0.00% | 0.56% | 0.63% | 0.00% | 0.00% | 0.14% | 0.11% | 0.02% | 0.00% |
| **Granulocyte** | 4.68% | 47.03% | 9.90% | 0.44% | 0.03% | 4.90% | 8.74% | 0.07% | 0.79% |
| **Granulocytopoietic** | 13.88% | 35.52% | 61.74% | 16.44% | 1.99% | 20.00% | 6.04% | 3.53% | 13.58% |
| **Hematopoietic** | 3.67% | 7.47% | 16.19% | 76.47% | 85.56% | 31.56% | 4.98% | 15.23% | 7.37% |
| **Macrophage** | 0.00% | 0.05% | 0.68% | 0.01% | 0.00% | 2.99% | 0.08% | 0.02% | 0.00% |
| **Monocyte** | 1.95% | 1.33% | 1.37% | 0.38% | 0.00% | 2.99% | 46.27% | 0.02% | 4.70% |
| **Proerythroblast** | 0.00% | 0.10% | 1.19% | 0.00% | 8.38% | 0.00% | 0.05% | 77.03% | 0.00% |
| **Promonocyte** | 3.12% | 1.73% | 3.41% | 3.77% | 1.24% | 10.07% | 31.93% | 1.59% | 73.16% |
| **T** | 0.47% | 0.18% | 2.80% | 0.44% | 0.08% | 1.50% | 0.33% | 0.00% | 0.01% |

**Table S1. Annotation and resolution of bone marrow cell types identified by scRNA-seq.** This table summarizes the annotation and reconciliation of bone marrow cell populations identified by single-cell RNA sequencing across all experimental groups. Unsupervised clustering was performed using Seurat, followed by reference-based cell type prediction using SingleR in combination with the Tabula Muris bone marrow reference dataset. The table reports the percentage of reference-predicted identities mapping to each Seurat cluster, and the final resolved cell type used for downstream analyses.
